# Environment-independent distribution of mutational effects emerges from microscopic epistasis

**DOI:** 10.1101/2023.11.18.567655

**Authors:** Sarah Ardell, Alena Martsul, Milo S. Johnson, Sergey Kryazhimskiy

**Author notes:** Equal contribution (Sarah Ardell, Alena Martsul). Platform Reagent Development Group, Illumina Inc., San Diego, 13 CA 92122.

## Abstract

Predicting how new mutations alter phenotypes is difficult because mutational effects vary across genotypes and environments. Recently discovered global epistasis, where the fitness effects of mutations scale with the fitness of the background genotype, can improve predictions, but how the environment modulates this scaling is unknown. We measured the fitness effects of ∼100 insertion mutations in 42 strains of *Saccharomyces cerevisiae* in six laboratory environments and found that the global-epistasis scaling is nearly invariant across environments. Instead, the environment tunes one global parameter, the background fitness at which most mutations switch sign. As a consequence, the distribution of mutational effects is predictable across genotypes and environments. Our results suggest that the effective dimensionality of genotype-to-phenotype maps across environments is surprisingly low.

**One Sentence Summary:** The effects of mutations on microbial growth rate follow a pattern of global epistasis that is invariant across environments.

## Main Text

Adaptive evolution can lead to profound changes in the phenotypes and behaviors of biological systems, sometimes with adverse and sometimes with beneficial consequences for human health, agriculture and industry (*1*–*4*). However, predicting these changes remains difficult (*5*–*7*). One major challenge is that how new mutations alter phenotypes and fitness of organisms often depends on the genetic background in which they arise (G×G interactions or “epistasis”), the environment (G×E interactions), or both (G×G×E interactions) (*8*). These interactions can alter not only the magnitude but also the sign of mutational effects, causing evolutionary trajectories to become contingent on the initial genotype, environment and random events (*9*).

Much of prior empirical work focused on characterizing “microscopic” G×G, G×E and G×G×E interactions for fitness-related phenotypes, that is, how the effects of individual mutations vary across genetic backgrounds and environments (*10*–*19*). Microscopic interactions determine evolutionary accessibility of mutational paths and the degree of genetic parallelism (*8, 10, 20*– *22*). However, using this approach for predicting genome evolution is challenging because the number of interactions grows super-exponentially with the number of variable loci (*7, 23*). Predicting phenotypic evolution may be more feasible and in many cases more useful (*7, 24*). Such predictions rely on coarse-grained, or “macroscopic”, descriptions of G×G, G×E and G×G×E interactions which ignore individual mutations and instead describe how the distributions of mutational effects vary across genotypes and environments (*8, 20, 25, 26*). While many studies measured the distributions of effects of mutations on fitness, or “DFEs” (*27*–*33*), a systematic understanding of macroscopic epistasis is still lacking (*26, 31, 33, 34*).

Several recent studies found that mutations often make the phenotype of an organism in which they occur less extreme (*11, 12, 34*–*42*), an instance of a more general phenomenon of microscopic “global epistasis” (*8, 43*). Such epistasis is expected to arise for complex traits, including fitness (*44*), and it can be used to quantitatively predict the effects of individual mutations in new genetic backgrounds without the full knowledge of their G×G interactions, thereby avoiding the combinatorial problem mentioned above (*43, 44*). Perhaps more importantly, if most mutations exhibit global epistasis, the distributions of their phenotypic effects should also have a quantitatively predictable shape (*44*), which could facilitate evolutionary predictions at the phenotypic level. However, we have limited understanding of how the patterns of microscopic global epistasis vary across environments (*19*), and no attempt has been made so far to empirically characterize how microscopic G×G, G×E and G×G×E interactions constrain the shape of the DFE.

Probing whether global epistasis models can capture G×G, G×E and G×G×E interactions at both microscopic and macroscopic levels hinges on measuring the effects of many mutations across multiple genetic backgrounds and environments. To this end, we measured how ∼100 quasi-random barcoded insertion mutations constructed in our previous study (*39*) affect growth rate in 42 “background” strains of yeast *Saccharomyces cerevisiae* in six conditions (*45*). All background genotypes are segregants from a cross between two strains of yeast (RM, a vineyard strain, and BY, a lab strain) and differ from each other by ∼2×10^4^ SNPs throughout the genome (*46*). Our environments varied by temperature (30°C and 37°C) and pH (3.2, 5.0 and 7.0), two stressors with global effects on yeast physiology (*47*–*49*), in a factorial design (see (*45*), Section 1.1 and Figure S1). This choice of strains and environments allowed us to explore a different (lower) range of growth rates of the background strains than in previous studies. Unlike previous studies, we kept our cultures growing close to the exponential steady state, which enabled us to infer the effect of each insertion mutation on absolute growth rate (denoted by λ) from bulk barcode-based competition experiments, with precision of about 7×10^−3^ h^−1^ (see (*45*), Section 1.3.3).

### Absence of unconditionally beneficial or deleterious mutations

We first estimated the effects of our mutations on growth rate in different background strains and environments. To this end, following Johnson et al (*39*), we designated a set of five mutations as a putatively neutral reference and found that the remaining 94 mutations exhibit a range of effects on growth rate relative to this reference (denoted by Δλ), from decreasing it by Δλ = – 0.18 h^−1^ to increasing it by Δλ = 0.14 h^−1^, with the median effect Δλ ≈ 0 h^−1^. We validated a subset of these estimates with an independent low-throughput competition assay (see (*45*), Section 1.2.3, Figure S2C). We then classified each mutation in each strain and environment as either beneficial or deleterious if the 99% confidence interval around its estimated effect did not overlap zero (see (*45*), Section 1.3.4). All other mutations were classified as neutral. This procedure yielded conservative calls of mutation sign, with a false discovery rate of 2.4%.

We found that the fraction of beneficial and deleterious mutations varied between 0% and 63% and between 0 and 59% per strain, respectively. The high proportions of beneficial mutations was unexpected, but not unprecedented (*30, 32*). However, no single mutation was identified as either beneficial or deleterious in all strains and environments. 94% (88/94) of our mutations are beneficial in at least one strain and condition, and, of those, 96% (85/88) are also deleterious in at least one strain and condition. Even within the same environment, between 33% (31/94) and 63% (59/94) of all mutations change sign across background strains, and between 1% and 41% of mutations change sign across environments in the same strain (Figure S3). Thus, the vast majority of mutations neither unconditionally increase nor unconditionally decrease growth rate.

A recent global epistasis model suggests that the fitness effects of most mutations should on average linearly decline with the fitness of the background genotype (*44*). One striking qualitative prediction of this model is that the proportion of beneficial mutations should increase with the decreasing fitness of the background strain, reaching up to 100% in the lowest-fitness genotypes, whereas the proportion of deleterious mutations should correspondingly decrease. Consistent with this prediction, we found that the proportion of beneficial and deleterious mutations increased and declined in slow-growing strains, respectively, and these relationships were statistically significant in 10/12 cases (Figure 1). Thus, global epistasis indeed appears to be a major determinant of the sign of mutations (Figure S3C).

**Figure 1.**
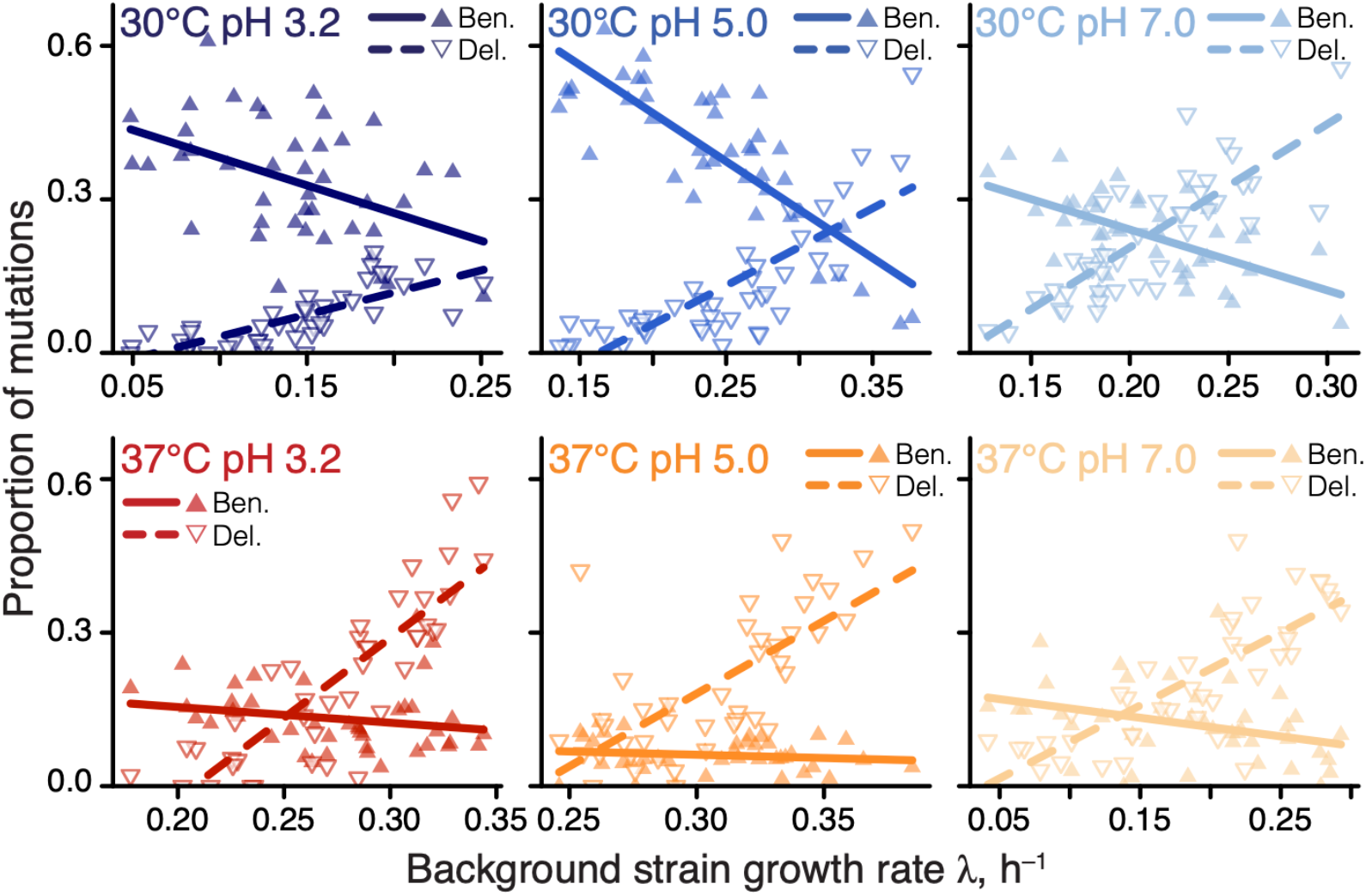
Proportions of beneficial and deleterious mutations vary with strain growth rate. Filled and empty triangles show the proportions of beneficial and deleterious mutations in each background strain as a function of its growth rate, respectively. Lines are the best-fit linear regressions; all are statistically significant (*P* < 0.05, t-test) except for beneficial mutations in 37°C pH 3.0 and 37°C pH 5.0.

### Consistency of microscopic global epistasis across environments

To probe the microscopic global epistasis model quantitatively, we modeled the effect Δλ_*mge*_ of each mutation *m* on growth rate in strain *g* and environment *e* as

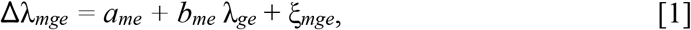

where λ_*ge*_ is the growth rate of the background genotype *g* in environment *e*. The first two terms in equation (1) capture global epistasis, a deterministic component which can be used for prediction, and ξ_*mge*_ captures the remaining (unpredictable) epistasis, which we refer to as “idiosyncratic” (*8, 36*).

We found that the linear model (1) was statistically significant for 97% (91/94) of mutations (*F*-test, *P* < 0.05 after Benjamini-Hochberg correction), and explained on average 45% (interquartile interval [30%, 61%]) of variance in the effects of mutations across background strains and environments (see (*45*) Section 1.3.7). When tested individually, 46% (250/545) of global epistasis slopes *b*_*me*_ are significantly different from zero (t-test, *P* < 0.05 after Benjamini-Hochberg correction), with 98% (245/250) of them being negative (Figure 2A), consistent with the global epistasis theory (*44*) and previous observations (*36, 39*). 96% (240/250) of significant intercepts *a*_*me*_ are positive (Figure 2B), implying that these mutations are expected to be beneficial in a hypothetical non-growing strain, consistent with relationships shown in Figure 1.

**Figure 2:**
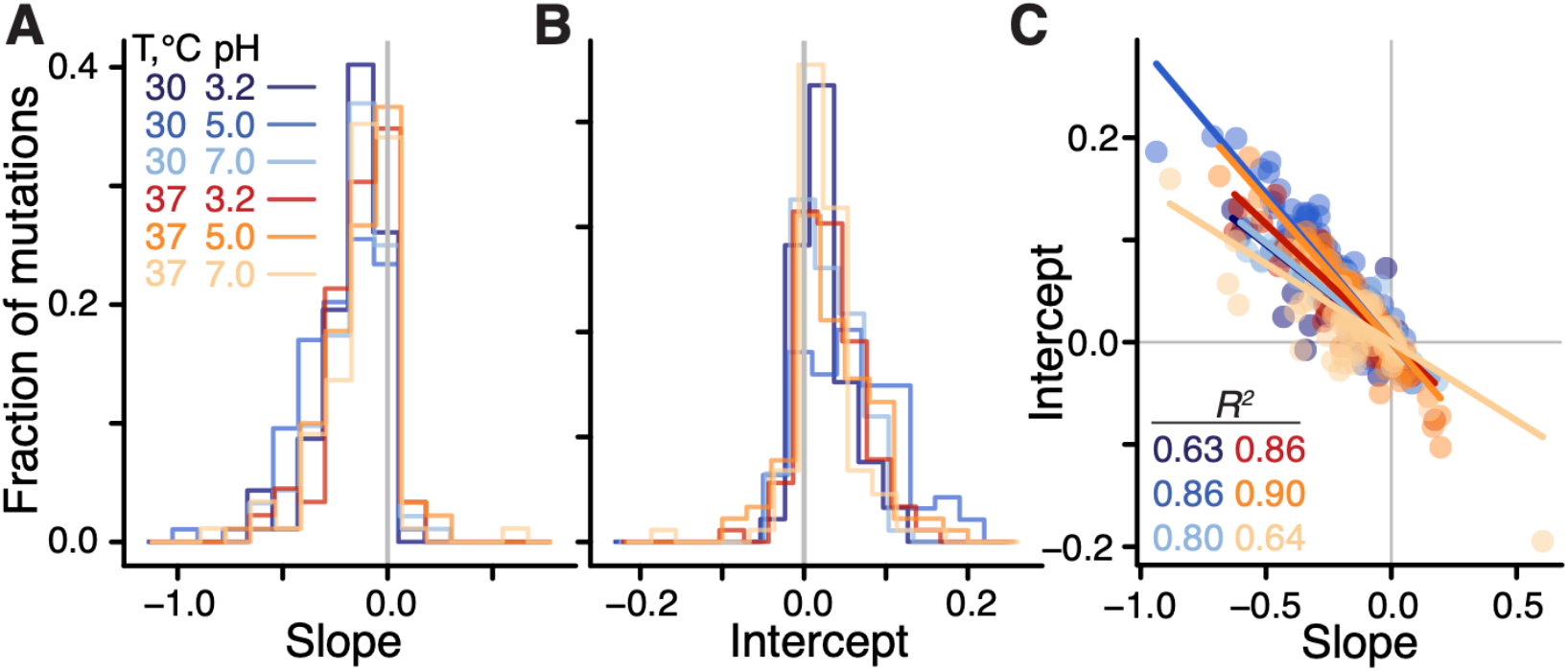
Distributions of global-epistasis slopes and intercepts and their correlation. A, B. Histograms of slopes and intercepts estimated from fitting equation (1) to data (see Figure S4 for statistical tests). **C**. Correlation between slopes and intercepts. Each point represents a mutation, colored by environment. Lines are the best fit linear regressions (*P* < 0.01 for all, t-test).

We next examined how the global epistasis slopes *b*_*me*_ and intercepts *a*_*me*_ vary across mutations and environments. We found that the distributions of slopes are nearly invariant across environments (Figures 2A, S4A,E and Table S1), while the distributions of intercepts are more variable (Figure 2B, S4A and Table S1). Furthermore, slopes and intercepts are strongly negatively correlated (Figure 2C, S4F), such that mutations with a zero slope have on average a zero intercept, which is consistent with the paucity of unconditionally deleterious and beneficial mutations noted above. Thus, global epistasis slopes and intercepts are mathematically related as 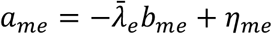 where the regression coefficient 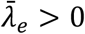 can be interpreted as the “pivot” growth rate at which a typical mutation switches its sign in environment *e* (see (*45*), Section 1.3.7). Each individual mutation switches its sign at a background growth rate that deviates from 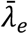 by η_*me*_/*b*_*me*_, and we therefore refer to η_*me*_ as the “pivot noise”. We find that the distributions of η_*me*_ in all environments are close to normal, with zero mean and standard deviation of approximately 1.7×10^−2^ (Figure S4C). On the other hand, pivot growth rates 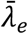 differ significantly across environments (see (*45*), Section 1.3.7; Figure S4 B,G).

There are two possibilities for how the distributions of global-epistasis slopes could be nearly invariant across environments. It could be because slopes of individual mutations are nearly invariant. Alternatively, slopes of individual mutations could change across environments, but their distribution is preserved. To test these hypotheses, we examined how the global-epistasis slope of each individual mutation changes across environments. We found that they are statistically indistinguishable in 87% (1169/1333) of pairwise comparisons, and 62% (58/94) of mutations have statistically indistinguishable slopes in all environments (Figures 3, S5, S6).

**Figure 3.**
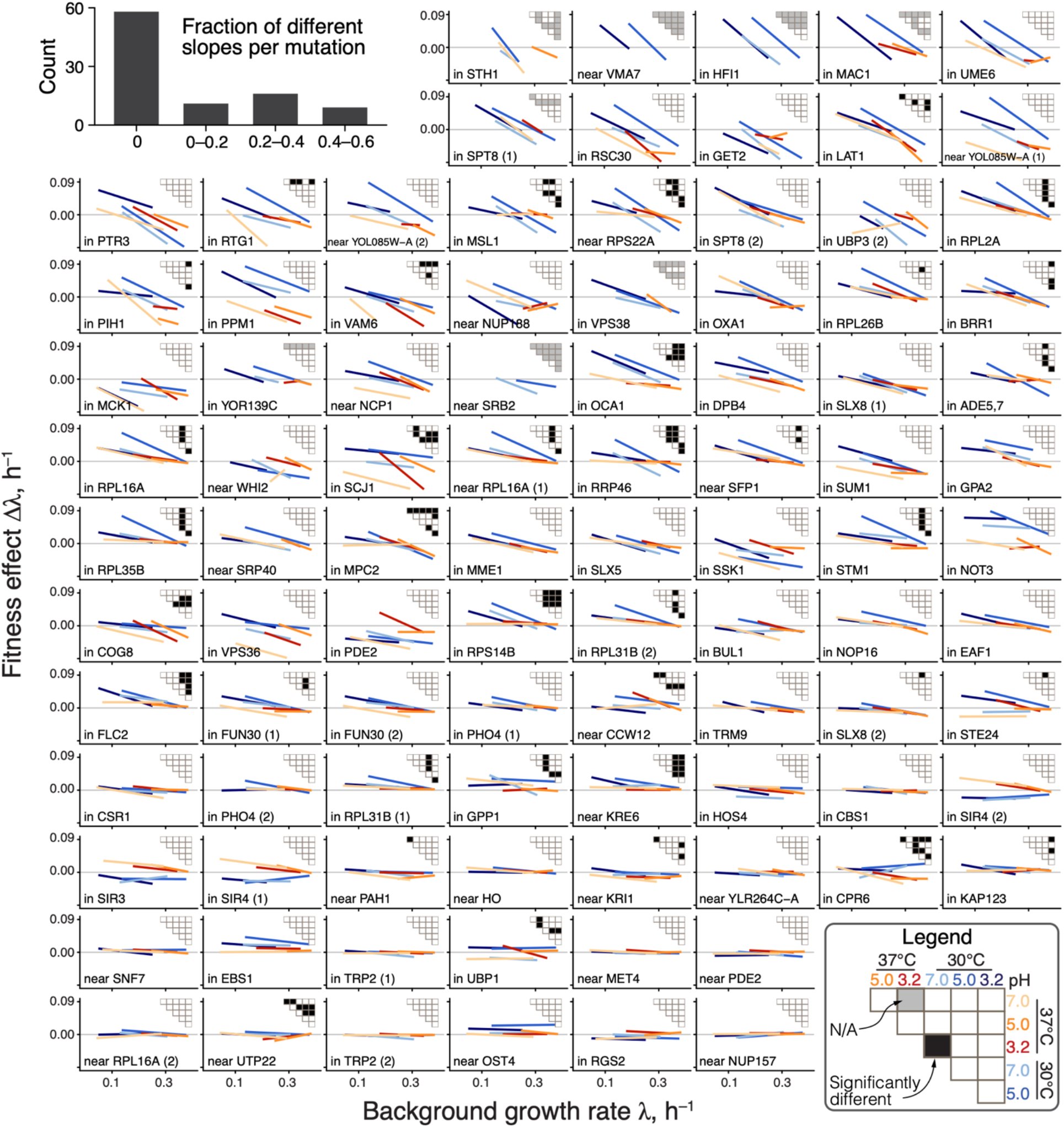
Global-epistasis slopes of mutations are nearly invariant across environments. Panels show regression lines from fitting equation (1) for each mutation, colored by the environment as in previous figures. Data points are shown in Figure S6. Mutations are displayed in the order of increasing mean slope. Insets show the results of all pairwise slope-comparison tests (legend in lower right). Histogram in top left shows the overall distribution of fractions of significant tests per mutation (see (*45*), Section 1.3.7).

Moreover, even when slopes are statistically distinguishable, they are very similar, so that a model with six environment-specific slopes *b*_*me*_ explains only 2% more variance in the effects of mutations compared to an “invariant slope” model where each mutation is characterized by a single environment-independent slope *b*_*m*_ (45% versus 43% on average; Figure S5). These results hold even if we only compare non-zero slopes or control for missing measurements (see (*45*), Section 1.3.7 and Figures S5).

The near-invariance of global-epistasis slopes of individual mutations could arise trivially if each environment shifted the growth rates of all strains by the same amount while preserving the relative order of their growth rates and the effects of mutations. However, this is not the case. We find that 34 out of 42 (81%) background strains are found among the 50% fastest growing strains in at least one environment and also among the 50% slowest growing strains in at least one other environment (see (*45*), Section 1.3.6 and Figure S7). Similarly, 50% of mutations on average switch between the top and bottom 50% of mutation effects in the same strain across environments (Figures S8). Thus, slopes are nearly preserved despite the fact that the relative rank orders of background strains and mutations are reshuffled across environments.

Taken together, the near-invariance of global-epistasis slopes across environments and the linear relationship between slopes and intercepts indicate that the microscopic G×G, G×E, and G×G×E interactions for most of our mutations follow a simplified version of equation (1),

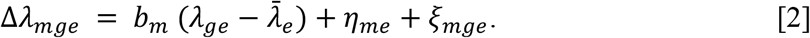

Equation (2), which we refer to as the “generalized global epistasis equation”, shows that the deterministic effect of the environment on global epistasis is captured by a single effective parameter, the pivot growth rate 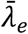.

### Distribution of fitness effects

To understand the implications of equation (2) for the macroscopic G×G, G×E and G×G×E interactions, we calculated the first three moments of the distribution of fitness effects of mutations (DFE) under the simplifying assumption that pivot noise η_*me*_ and idiosyncratic epistasis ξ_*mge*_ terms are all independent (see (*45*), Section 1.4). The generalized global epistasis equation predicts that the DFE mean ought to decline linearly with the background growth rate, with the same slope across environments, crossing zero at the environment-specific pivot growth rate 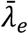.

The behavior of higher DFE moments is less obvious. DFE variance is predicted to depend on λ_*ge*_ quadratically, with the parabola’s minimum achieved at the pivot growth rate. To understand this prediction intuitively, consider an idealized case without idiosyncratic epistasis ξ_*mge*_ = 0 (Figure 4A). Since most mutations switch sign near the pivot growth rate, strains growing at this rate have access only to mutations with small effects, and the DFE is narrow. As the background growth rate deviates from 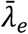, global epistasis lines for individual mutations spread out forming a “bowtie” (Figure 4A), and the DFE variance increases in both directions. This general pattern still holds even when ξ_*mge*_ ≠ 0 (see (*45*), Section 1.4).

**Figure 4.**
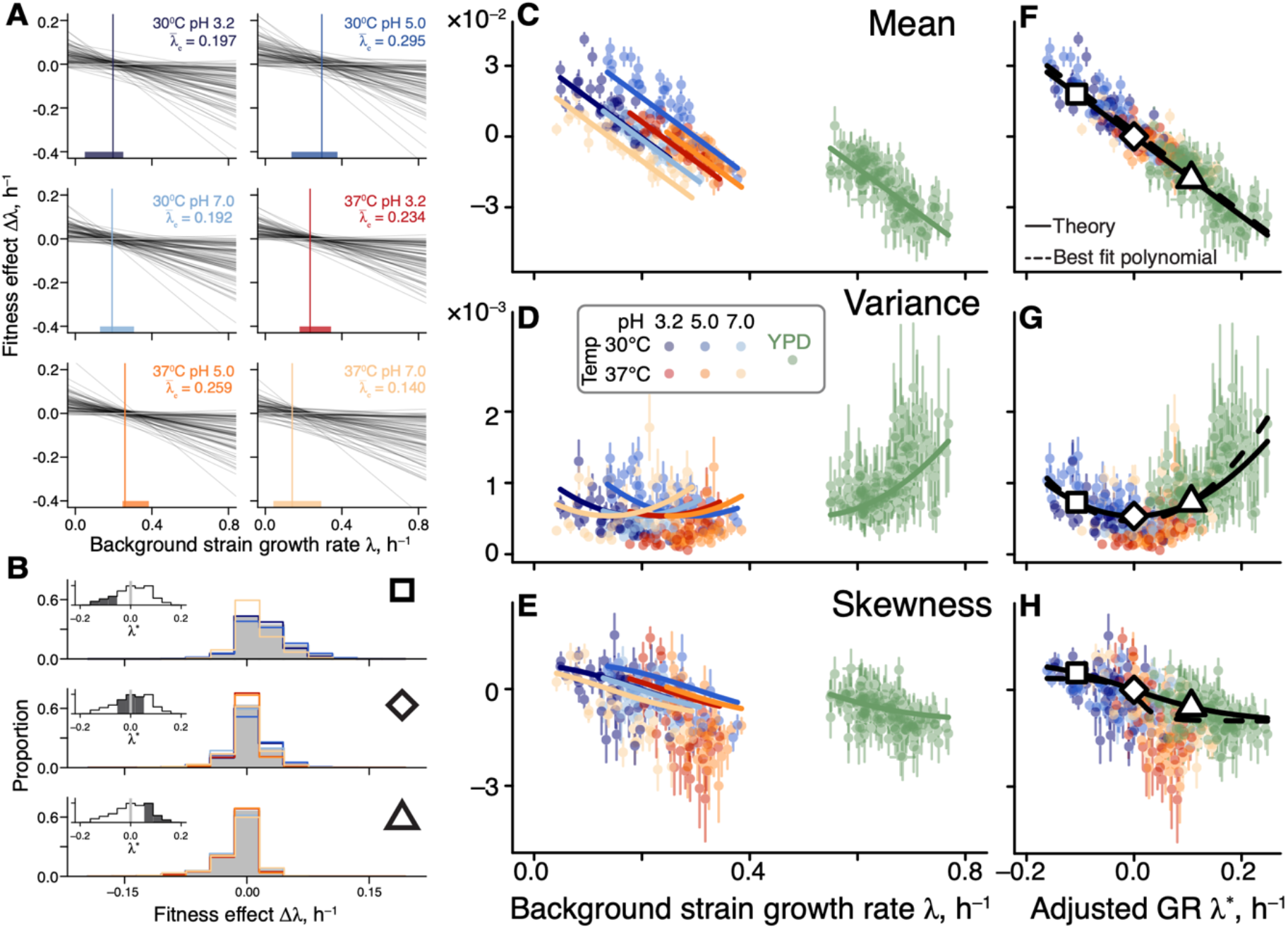
Generalized global epistasis equation captures changes in the DFE across strains and environments. **A**. The global epistasis contribution to the fitness effect of each mutation (term 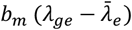 in equation (2)) as a function of background growth rate term λ_*ge*_. Each line represents a mutation. Vertical line indicates the pivot growth rate. Thick colored lines on the *x*-axis indicate the range of measured background-strain growth rates in each environment. **B**. Estimated DFEs for strains whose adjusted growth rate is negative (top panel), approximately zero (middle panel) and positive (bottom panel). Gray bars show DFEs pooled across all environments, colored lines show DFEs for individual environments (colors are as in previous figures). Insets show the distributions of adjusted growth rates for background strains, with the focal bin shaded. Large square, rhombus and triangle are shown for reference with panels F,G,H. **C, D, E**. DFE moments plotted against the background strain growth rate. Error bars show ±1 standard errors (see (*45*), Section 1.3.9). Solid curves show the theoretical predictions calculated from equation (2) and parameterized without the YPD data (see (*45*), Section 1.4). DFE moments are calculated with a median of 74 mutations (interquartile interval [61,79]). **F, G, H**. Same data as in C, D, E, but plotted against the adjusted growth rate. Dashed curves are the best fitting polynomial of the corresponding degree (see (*45*), Section 1.4).

Finally, DFE skewness is predicted to decline monotonically with λ_*ge*_, crossing zero again at the pivot growth rate. More generally, our model makes a qualitative prediction that the DFE varies across strains as a function of their environment-adjusted growth rate 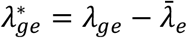 rather than as a function of their absolute growth rate λ_*ge*_. Furthermore, when the adjusted growth rate is zero, all odd central moments of the DFE are predicted to vanish and all even central moments are predicted to achieve their minimum (see (*45*), Section 1.4).

To test these predictions, we compared the empirical DFEs across environments. We find that pairs of strains with matched adjusted growth rates have significantly more similar DFEs than pairs of strains with the same absolute growth rate in different environments or the same strain in different environments (Figure 4B and S9), consistent with our predictions. We then plotted the first three moments of the empirical DFEs against the unadjusted and adjusted growth rate in all environments. We find that these moments align well only in the latter plot (Figures 4C–H). As predicted, the DFE mean and skewness decline monotonically with the strain’s adjusted growth rate 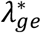 and cross zero when 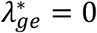, and DFE variance achieves its minimum at 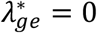. We confirmed that these patterns are not a result of missing measurements (see (*45*), Section 1.3.9 and Figure S10).

The fact that all of our predictions hold indicates that the linear generalized global epistasis equation with uncorrelated noise terms quantitatively captures a major mode of variation in the DFE shape across genotypes and environments. However, if the environment truly modulates only one effective parameter, the pivot growth rate, then we should be able to predict DFE shapes in any environment, once its pivot growth rate is known. To test this prediction, we turned to our previous work where we measured DFEs of 163 yeast strains (a superset of the 42 strains used in this study) in a rich medium YPD (*39*). We estimated the pivot growth rate for this environment as the background growth rate where the DFE mean equals zero (see (*45*), Section 1.3.5). After growth-rate adjustment, we found that our theoretical predictions quantitatively capture variation in the DFE variance and skewness without any other fitted parameters (green points in Figure 4C-H).

## Discussion

Having measured the fitness effects of many mutations across diverse yeast genotypes and environments, we obtained two main results. First, the global-epistasis slopes of individual mutations are nearly invariant across environments, which leads to a simple equation (2) that predicts the expected fitness effect of a mutation in a new strain and environment. Following Reddy and Desai (*44*), we interpret this equation as a statistical description of an underlying canonical (i.e., deterministic) genotype-to-fitness map. In fact, it is straightforward to find genotype-to-fitness maps that quantitatively reproduce the patterns of global epistasis that we observe in our data (see (*45*), Section 2.3). Within the Reddy-Desai model, the global
-epistasis slope of a mutation at a given locus is related to the fraction of variance in fitness attributable to this locus. Thus, the near-invariance of global-epistasis slopes suggests that our environmental perturbations largely preserve the overall statistical structure of genetic interactions, consistent with previous work in yeast (*50*). Whether this regularity holds for more extreme perturbations or those that target specific cellular processes, such as exposure to drugs, remains to be seen (*19*).

Our second main result is the empirical characterization of constraints imposed on the DFE by microscopic global epistasis. We found that the environment controls a single effective parameter, the pivot growth rate 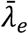, and a genotype’s DFE is set by the deviation of its growth rate from 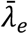, as shown in Figure 4. For example, many of our background strains happen to grow slower than 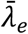 and, as a result, have access to many beneficial mutations (Figure 1). The physiological mechanisms that determine the pivot growth rate are as of yet unclear. According to the Reddy-Desai model, “bowtie” patterns analogous to those shown in Figure 4A are expected to arise generically on genotype-to-fitness maps with many epistatic interactions, essentially as a consequence of the regression to the mean (*44*). In this model, a typical mutation switches sign at the average fitness of all genotypes, indicating that the pivot growth rate could be a metric of environmental permissibility. The fact that in our data the pivot growth rate correlates with the average growth rate of our background strains is consistent with this interpretation (Figure S4G).

Overall, our results bolster the Reddy-Desai theory. However, whether this theory provides the most accurate explanation for global epistasis remains to be determined. One potential shortcoming of this theory is that it ignores the biological architecture of organisms, which could impose some non-trivial structure on epistasis (*51*).

Regardless of the biological mechanisms that underlie global epistasis, further work is needed to establish its generality. If it holds broadly, several important implications are conceivable. In genetics, our model can improve predictions of the phenotypic effects of mutations. In conservation biology, the possibility that low-fitness genotypes have access to large supplies of beneficial mutations gives hope that evolutionary rescue may prevent some species extinctions. In evolutionary biology, our results point to the existence of a universal class of distributions of fitness effects of mutations, which could explain why evolutionary dynamics of fitness are so similar and predictable across systems (*25, 36, 42, 52*). More broadly, our model may help us better understand these dynamics, including how the DFE changes during evolution (see (*45*), Section 2.2), how adaptation to one environment affects fitness in another (*26*), how pan-genomes evolve (see (*45*), Section 2.3), etc. More fundamentally, our results suggest that epistasis reduces the effective dimensionality of genotype-to-phenotype maps (*8, 43, 44, 51*). What biological constraints cause this dimensionality reduction, how it emerges and when it breaks down are exciting open questions in systems biology.

## Acknowledgements

We thank the members of the Kryazhimskiy and Meyer labs, Michael Desai, Terry Hwa and Erik Hom for insightful discussions, Gautam Reddy, and three reviewers for constructive feedback on the manuscript. Computational work was assisted by the UC San Diego Triton Computing Cluster.

## Author contributions

Conceptualization: SMA, SK; experimental design: AM, MSJ, SK; experiments: AM; analysis: SMA, SK; writing: SMA, MSJ, SK.

## Data and materials availability

Data described in the paper are presented in the supplementary materials (Tables S1 through S6). Raw sequencing data are publicly available at the NCBI Sequence Read Archive (accession no. PRJNA1028648), and all analysis code is available on Zenodo (*53*). Data for the YPD environment provided in Ref. (*39*).

## Supplementary Materials

Materials and Methods

Supplementary Text

Figs. S1 to S13

Tables S1 to S4

References (54–71)

## Supplementary Materials for

### 1 Materials and methods

#### 1.1 Strains and media

##### 1.1.1 Background strains

Our “background strains” of yeast *S. cerevisiae* are a subset of a larger library of segregants that were previously generated from a cross between the lab strain BY and the vineyard strain RM (*46*) and whose evolutionary properties have been previously characterized (*34, 39, 54*). Specifically, our set of 42 background strains (listed in Table S6-Tab 2) is a subset of the strains used in the “Small Library” RB-TnSeq experiment described in Ref. (*39*). All the necessary strain details can be found in Refs. (*39, 46*). Most importantly, our background strains differ from each other at approximately 25,000 loci and span nearly the full growth rate range in YPD measured in Ref. (*39*) (see Section 1.3.5). These strains also vary widely in both DFE mean (2.5-fold range) and variance (1.5-fold range) in YPD.

##### 1.1.2 RB-TnSeq libraries

Background strains individually transformed with these RB-TnSeq libraries were kindly provided by Michael Desai (Harvard University). The design and construction of the RB-TnSeq libraries is described in detail in Ref. (*39*). Briefly, each background strain was transformed twice with the same set of 100 redundently barcoded transposon-insertion mutations, resulting in two biological replicates for each mutation in each strain, such that, within each transformation, each barcode uniquely tags a particular mutation and background strain (43% of barcodes were used in both transformations). This set of mutations is conditionally random, given two constrains, as described in Ref. (*39*). Briefly, (i) mutations whose fitness effects we can measure must be non-lethal and (ii) only those mutations were included in this set that had a measurable fitness effect in at least six out of 18 genetic backgrounds in the rich medium YPD. Thus, the results of this work should be viewed as representing trends in the effects of disruptions of non-essential but functionally important genes.

On average, each mutation was represented by 11 and 37 barcodes in the first and second transformation, respectively (see Table S6-Tab 5). Five mutations (IDs: 91 (nearby COA6), 51 (nearby FIT2), 6 (nearby MET2), 99 (nearby TDA11), 102 (nearby YSP2)) that target intergenic regions were used as a neutral reference, as in Ref. (*39*). These mutations were found to have no signficant fitness effects in all strains in one environment in Ref. (*39*). These reference mutations were represented by on average 19 and 77 barcodes in the two transformations.

##### 1.1.3 Media and environments

Unless otherwise noted, all experiments were performed in synthetic complete medium (SC, 2% dextrose (VWR, #90000-904), 0.67% YNB + nitrogen powder (Sunrise Science Products, #1501-500), 0.2% synthetic complete drop-out powder mixture (Sunrise Science Products,#1300-030)). We added ampicillin (Amp) and tetracycline (Tet) at concentrations given in Table S2 into the medium to prevent bacterial contamination. Our environmental conditions differed by two factors, temperature (30°C and 37°C) and pH (3.2, 5.0, and 7.0). pH was maintained with the citrate-phosphate buffer (*55*), which was prepared using 1 M stocks of citric acid (VWR, # 97061-858) and K_2_HPO_4_ (VWR, #97062-234) following the protocol described in Ref. (*56*). Autoclaved media were pH-adjusted by adding the necessary volumes of sterile citric acid and K_2_HPO_4_ solutions and measuring the pH of the buffered media. If the pH of the buffered media deviated from the desired level, we further adjusted it by adding small volumes of 4 M HCl (Sigma-Aldrich #84435). Media was used within two days.

#### 1.2 Experimental procedures

##### 1.2.1 RB-TnSeq experiment

To estimate the effects of tn-mutations on the absolute growth rate in 42 background strains in 6 environments, we pooled the tn-mutant libraries of our background strains into multiple pools (see below for details) and maintained these pools in continuous growth in 150 ml of media in 500 ml flasks over the period of 48 hours with dilutions down to 5 *×* 10^7^ cells and sampling every 12 hours. We carried out two replicate competition assays, one per independently transformed library (see Section 1.1.2).

###### Pre-growth and pooling

Prior to pooling, mutagenized strain libraries were defrosted and pre-grown for 24 h in 96-deep-well plates with 1 ml of media in each of our environments. Based on preliminary growth rate estimates, we grouped the background-strain libraries by their growth rate into three groups for the competition assays at 30°C and into two groups for the competition assays at 37°C. Each background strain was represented only in one group per temperature, with the exception of LK5-G01, which was added to each group as cross-group control. Group identities of each background strain can be found in Table S6-Tab 2. Thus, we propagated 6 cultures (3 growth rate groups × 2 biological replicates) in each of the 30°C environments and 4 cultures (2 growth rate groups × 2 biological replicates) in each of the 37°C environments for a total of 30 cultures.

After pre-growth, we measured OD600 of each mutant culture and converted it into cell density using a previously obtained calibration curve. Based on these density estimates, we pooled mutant cultures at approximately equal abundances, with a slight over-representation of those background strains whose preliminary growth rate estimates were lower. We then measured the density of each mixed culture again and transferred 5 *×* 10^7^ cells into the corresponding competition flask. The remaining mixed *T*_0_ cultures were frozen using the protocol described below.

###### Growth and dilution

Competitions were carried out in 150 ml of media in 500 ml baffled flasks (Pyrex No. 4446-500) in a shaking incubator (Eppendorf New Brunswick I26, 2.5 cm orbit) set to 150 rpm and appropriate temperature. Every 12 h, we estimated the cell density of each culture using plate-reader-based OD600 measurements and a previously obtained calibration curve (Table S6-Tab 3). Then, 5 *×* 10^7^ cells were transferred into the fresh media. When the transfer volume exceeded 1 ml, we adjusted the volume of fresh media to maintain the consistent culture volume of 150 ml. All cultures were propagated for four growth and dilution cycles (48 hours), yielding five samples per culture.

###### Sampling and storage

We pelleted the cells from 50 ml of each cultures remaining after the transfer and froze the cell pellets at *−*70°C for subsequent DNA extraction and barcode sequencing.

##### 1.2.2 Sequencing library preparation

We used the YeaStar Genomic Kit Protocol I (Zymo Research, #D2002) to extract gDNA from *∼* 1 ml of pelleted yeast cultures. To generate Illumina-ready dual-indexed amplicon library, we used a two-step PCR protocol, modified from Ref. (*29*) as follows. All primer sequences can be found in Table S5.

###### First PCR

We combined 100 ng of extracted gDNA, 25 µl of OneTaq DNA polymerase Master Mix (New England BioLabs, #M0482L), 0.5 µl of 10 µM oAM-R2P-100-R01 primer, µl of 10 µM oAM-R1P-20X-F01 primer, 1 µl of 50 µM MgCl_2_, and molecular biology grade water up to the total volume of 50 µl. We used the following PCR protocol:

1. 94°C for 30 sec.
2. 94°C for 30 sec.
3. 50.5°C for 30 sec.
4. 68°C for 70 sec.
5. Repeat Steps 2–4 for a total of three times
6. 68°C for 5 min.

We purified this PCR product with AMPure XP magnetic beads (Beckman, #A36881) (1:1 ratio).

###### Second PCR

We combined 15 µl of purified PCR I product, 25 µl of OneTaq DNA polymerase Master Mix, 1 µl of 50 µM MgCl2, 1 µl of 10 µM N7XX primer (Nextera), and 1 µl of 10 µM S5XX primer (Nextera), and 7 µl of molecular biology grade H2O. We used the following PCR protocol:

1. 94°C for 30 sec.
2. 94°C for 30 sec.
3. 62°C for 30 sec.
4. 68°C for 70 sec.
5. Repeat Steps 2–4 for a total of 24 times
6. 68°C for 5 min.

The final PCR product was purified as above, run on a gel, extracted and purified with QIAquick PCR purification kit (Qiagen, #28106).

###### Sequencing

We sequenced the libraries with paired-end 150 bp reads on one HiSeq4000 platform and two HiSeq X10 platforms (Illumina).

##### 1.2.3 Validation of estimated fitness effect of mutations

We were surprised by the high fraction of beneficial mutations identified in our RB-TnSeq experiment and carried out an additional experiment designed to validate our fitness-effect estimates obtained in the large-scale multi-strain pooled experiment. Specifically, we wanted to validate that the fitness effects of mutations (i) were in fact driven by the tn-insertions themselves rather than some additional unobserved mutations that could have arisen during transformation, and (ii) were not too specific to the environment created in the multi-strain competition pools.

To this end, after preliminary analyses, we selected a set of seven mutations that were expected to span a range of fitness effects: nearby MET2, nearby MET4, in NOT3, in PPM1, in RSC30, nearby TDA11, in MPC2 (Mutation IDs: 6,10, 117, 66, 71, 99, and 127, Table S6-Tab 4). We generated new RB-TnSeq libraries for these mutations and reconstructed them in five background strains, LK2-D07, LK6-A05, LK5-C04, LK1-H02, and LK1-C09. Two of the selected mutations (nearby MET2 and nearby TDA11, IDs: 6, 99) were used as a neutral reference in the RB-TnSeq experiment. The mutation nearby MET4 (ID: 10) was identified as neutral in the backgrounds and environments used in the validation experiment, and the remaining mutations were identified as non-neutral in at least some of these strains and environments.

The barcoded libraries for individual mutations were generated using the protocol described in Ref. (*39*). In total, we created 28 barcoded plasmid libraries (two replicated libraries per mutation, and two different drug resistance markers per mutation and replicate), such that each library contained a unique set of barcodes. We then transformed these libraries into all background strains, with LK2-D07 and LK6-A05 receiving one drug resistant marker (Hygromycin) and the remaining three LK5-C04, LK1-H02, and LK1-C09 receiving the other (Nourseothricin), again following protocols described in Ref. (*39*). Transformant colonies were scraped and, after 24 h of additional growth in selective media, cultures were pelleted and frozen in 20% glycerol at *−*70°C. To determine which barcodes were associated with each mutation in each background strain, we sequenced each of the 28 yeast mutant libraries at the barcode locus using the same protocols as in Section 1.2.1.

After generating the libraries of individual mutations, we estimated their fitness effects in two environments, 30°C pH 5.0 and 37°C pH 7.0, by competing only mutants that share the same genetic background. We pre-grew 140 cultures (5 background strains *×* 7 mutations *×* 2 replicate mutant libraries *×* 2 environments) in the buffered SC media containing Amp (to avoid bacterial contamination) and either Nat or Hyg (required for selecting for the transformants, see Tables S6-Tab 2 and S2, and Ref. (*39*)) and incubated them in the two focal environments (30°C pH 5.0 and 37°C pH 7.0) for 24 h in test tubes shaken at 220 rpm. After measuring the concentrations of the grown cultures as described above (see Section 1.2.1), we created one mixed culture per background strain, per mutant library and per environment (2 environments *×* 5 strains *×* 2 biological replicates, for a total of 20 mixed cultures) as follows. We added each of the two neutral reference mutation cultures (IDs: 6,99) at frequency 25% each and we added each of the “query” mutant cultures (IDs: 10, 117, 66, 71, and 127) at 10%, such that initial ratio of reference and query mutants in each culture was 1:1. After estimating cell counts in these mixed cultures using OD600 (see Section 1.2.1), we transferred 5 *×* 10^7^ cells to start the competition assay. The assays, sampling and sequencing library preparation were performed following the same protocols as in Section 1.2.1. Fitness effects of mutations estimated in this experiment are provide in Table S6-Tab 8.

#### 1.3 Data Analysis

##### 1.3.1 Code

All analysis code is written in R 4.3.1 and is available at https://github.com/ardellsarah/Yeast mutation effects across strains and environments. and archived at DOI 10.5281/zenodo.12696517.

Packages used are listed in the beginning of all scripts. Computationally intensive analyses were run on the Triton Supercomputing Cluster (TSCC).

##### 1.3.2 Counting barcodes

###### Raw barcode counts

Barcode counts for the main RB-TnSeq experiment (Section 1.2.1) were obtained using the BarcodeCounter2 package (*57*) with a pre-determined barcode-mutation association data from Ref. (*39*).

To determine barcode counts in the validation experiment, we first used the barcode sequencing data for individual mutation libraries (see Section 1.2.3) to associate each barcode sequence with a particular mutation and background strain. To do so, we used regular expressions to extract all unique barcode sequences and clustered them using the `seq cluster` function in R’s bioseq package. We then used BarcodeCounter2 with the resulting barcode-mutation associations to extract raw barcode counts for each sample file.

###### Filtering

Because it is critical to have accurate reference barcode counts for the inference of fitness effects of mutations, we discarded all time points that contained less than 500 reference mutation counts for any given strain, replicate and environment. This filtering removed 1.3% (41,404/3,060,514) of strain-environment-replicate-time point combinations. Then, we retained only those barcodes that were present at three or more time points (in any given condition and replicate) at 5 or more counts at each time point.

##### 1.3.3 Estimating growth rates of background strains and fitness effects of mutations

The central piece of our procedure for estimating the growth rates of background strains and the fitness effects of mutations is the detection of “outlier” barcodes, i.e., those that have abnormally high or low growth rates relative to other barcodes tagging the same mutation. Such outliers arise likely due to the tn-mutants acquiring secondary mutations either during the barcoding step or during the RB-TnSeq experiment. The outlier detection procedure requires preliminary estimates of fitness effects of barcodes tagging each mutation, which in turn requires a robust set of reference barcodes. Thus, our procedure consists of the following steps.

1. Estimate the growth rates of all barcodes.
2. Detect and exclude outlier barcodes for reference mutations to obtain a robust estimate of the growth rate of reference mutants.
3. Obtain preliminary estimates of fitness effects of mutations based on the robust set of reference barcodes.
4. Detect and exclude outlier barcodes for non-reference mutations.
5. Estimate the background growth rates and the fitness effects of mutations in each biological replicate.
6. Pool estimates across replicates.

We describe the outlier detection algorithm at the end of this section. Suffices to say that this algorithm takes as input (i) the set of all barcodes tagging a given mutation (in a given genetic background, biological replicate and environment), (ii) the cell count estimates over time of all these barcoded lineages and (iii) the preliminary estimates of the fitness effects of these lineages, and it outputs a robust “outlier-free” set of barcodes corresponding to the mutation.

Before we describe our estimation procedure, recall that, in a given biological replicate *r*, each barcode *k* uniquely specifies a particular tn-mutation *m* and the background strain *g* into which this mutation is introduced (see Section 1.1.2). We denote the set of all barcodes tagging mutation *m* in genetic background *g* in replicate *r* by 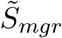. We also denote the set of five reference mutations by 𝒮^ref^ (see Section 1.1.2 and Table S6-Tab 4) and we denote the set of “reference barcodes”, i.e., all barcodes that tag these reference mutations in the genetic background *g* and replicate *r*, by 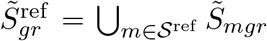. Tilde denotes the fact that these sets potentially include outlier barcodes.

###### Estimation of barcode growth rates across time intervals

We first calculate the frequency of each barcode *k* (reference or non-reference) in each replicate *r* at each time point *t* by dividing its read count by the total count of all barcodes present in the same flask at that time. To estimate the number of cells *N*_*kret*_ that carry barcode *k* in repliate *r* in environment *e* at the sampling time *t*, we multiply barcode frequency by the total number of cells present in the flask at that time point (Table S6-Tab 3). We estimate the growth rate *λ*_*kret*_ of the barcode lineage *k* as

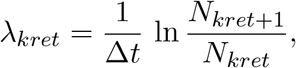

where Δ*t* is the time between transfers (typically 12 h, Table S6-Tab 3).

###### Detection and exclusion of outlier reference barcodes

We obtain a preliminary estimate of the effect of each reference barcode *k* in background *g*, environment *e* and replicate *r* as

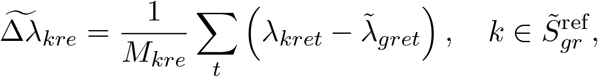

where 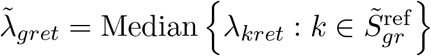 and *M*_*kre*_ is the number of time intervals where barcode *k* is observed in environment *e* in replicate *r*. We use these estimates to apply our outlier detection algorithm (see below) and generate a robust set of reference barcodes 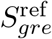 for each background strain *g* in each environment *e* and biological replicate *r*.

###### Preliminary estimates of fitness effects of mutations

For any barcode *k* tagging a non-reference mutation *m* in genotype *g*, we calculate its fitness effect in environment *e* and replicate *r* as

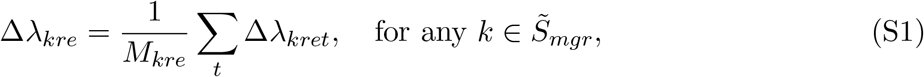

where *M*_*kre*_ is the number of time intervals when this barcode is observed in environment *e* in replicate *r*, and

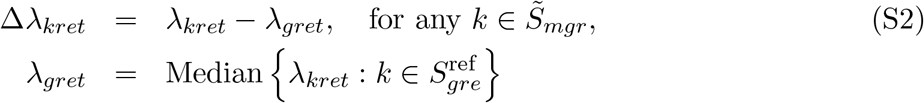

are robust estimates of the fitness effect of barcode *k* and growth rate of the background strain *g*, respectively, both at time interval *t* in environment *e* and replicate *r*. We then obtain a preliminary estimate of the fitness effect of each mutation *m* in genetic background *g* in environment *e* and biological replicate *r* as

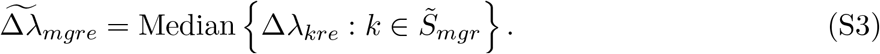

###### Detection and exclusion of outlier barcodes for non-reference mutations

We use preliminary fitness effect estimates given by equation (S3) and apply our outlier detection algorithm (see below) to generate a clean set of barcodes *S*_*mgre*_ for each mutation *m* in each background strain *g* in each environment *e* and biological replicate *r*.

###### Estimation of background growth rates and fitness effects of mutations in each biological replicate

We estimate the growth rate of each background strain *g* in each environment *e* and biological replicate *r* as

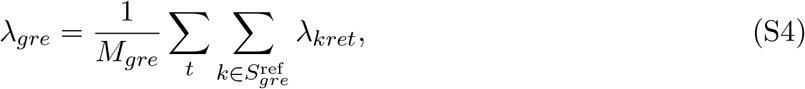

where *M*_*gre*_ is the number of barcode-time interval combinations at which *λ*_*kret*_ are estimated.

We estimate the fitness effect of each mutation *m* in each genetic background *g* environment *e* and biological replicate *r* as

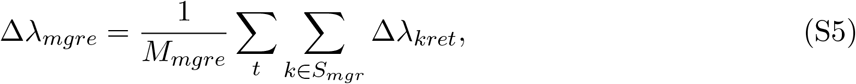

where Δ*λ*_*kret*_ are given by equation (S2) and *M*_*mgre*_ is the number of barcode-time interval combinations at which *λ*_*kret*_ are estimated.

###### Pooling estimates across neutral references

We estimate how the growth rate estimates of background strains correlate between estimates from single references and all four other reference mutations combined. We find they are highly correlated (Figure S2F), so we pool the data from all reference mutations together to obtain final growth rate estimates of each background.

###### Pooling estimates across biological replicates

We estimate how the fitness effects of mutations obtained using equation (S5) correlate across replicates. Since the replicates are highly correlated (Figure S2A-B), we pool the data from both replicates to obtain our final estimates of background growth rates and their standard errors,

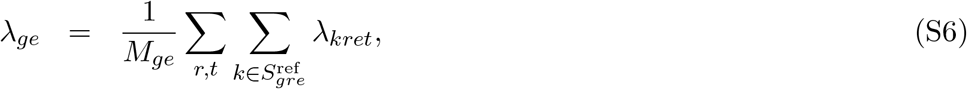

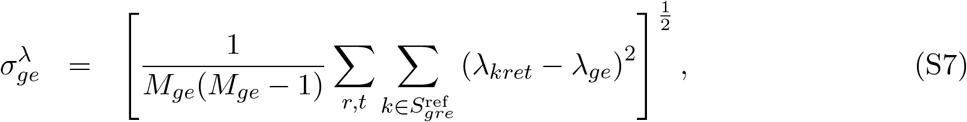

where *M*_*ge*_ is the number of barcode-replicate-time combinations at which *λ*_*kret*_ are estimated for background *g* in environment *e*. We estimate the fitness effects of mutations and their standard errors

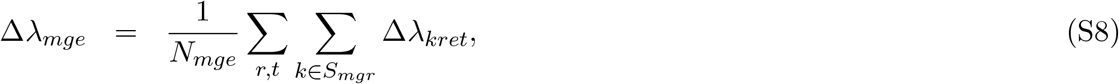

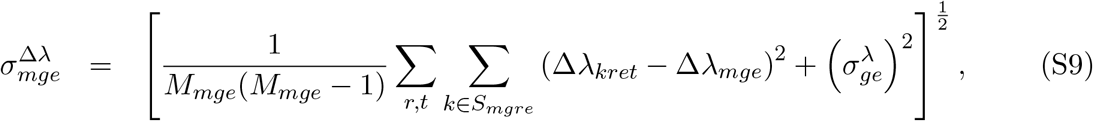

where *M*_*mge*_ is the number of barcode-replicate-time combinations at which Δ*λ*_*kret*_ are estimated for mutation *m* in background strain *g* and environment *e*.

The distributions of standard errors for background growth rates and fitness effects are shown in Figure S2G. The average standard error of the background growth rate is

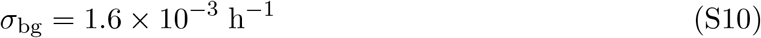

and the average standard error of fitness effect is

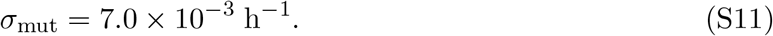

The comparison of these values shows that the noise in non-reference barcodes is typically more than 4-fold higher than noise in reference barcodes.

###### Detection of outlier barcodes

The goal of this procedure is to detect those barcodes whose frequencies either rise or fall unexpectedly quickly compared to the other barcodes tagging the same mutation in the same genetic background. To this end, we follow the method developed in Ref. (*39*), which takes as input the set of all barcodes *k* tagging a given mutation (in a given genetic background, biological replicate and environment), the corresponding cell counts *N*_*kt*_ at each time sampling *t* and the preliminary estimates 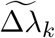 of the fitness effects of all these barcoded lineages. Briefly, we calculate the “within-mutation” frequencies *f*_*kt*_ at time *t* as 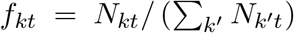. All barcoded lineages tagging the same mutation should grow at the same rate. Therefore, we expect all frequencies *f*_*kt*_ to be constant over time, barring demographic and sampling noise. To determine which frequency trajectories are inconsistent with this neutral expectation, we create a “within-mutation-neutral reference” (WMNR) set of barcodes whose preliminary fitness effect 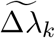 is within 0.01 of the median fitness effect of all these barcodes. Then, we fit two models to all frequency trajectories *f*_*kt*_ for those barcodes that are not in the WMNR set. In the neutral model, each query barcode’s trajectory does not systematically change relative to the pooled WMNR trajectory. In the model with selection, query barcode’s frequency can systematically increase or decrease. We find the log-likelihood of the observed barcode trajectory given each of the two models and calculate the likelihood ratio (LR) statistic. We conservatively exclude all barcodes with the LR statistic values greater than 40, corresponding to a *P* -value from a *χ*^2^ distribution with 1 d.f. *<* 10^−9^. This algorithm returns a “clean” subset of barcodes that tag a given mutation (in a given genetic background, environment and replicate).

Using this method, we exclude a total of 2.4% (16,799/683,754) of barcode-replicate combiations.

##### 1.3.4 Calling beneficial and deleterious mutations

To call each mutation as either beneficial, deleterious or neutral in each genetic background and environment, we construct the 99% confidence interval using a normal distribution with the mean equal to Δ*λ*_*mge*_ (equation (S8)) and variance equal to the standard error of the mean 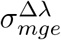 (equation (S9)). Mutations whose entire confidence interval is below zero are called deleterious and those whose entire confidence interval is above zero are called beneficial. All other mutations, i.e., those whose confidence interval spans zero, are called neutral. We identify a total of 6971 non-neutral mutation-genotype-environment combinations out of 17306 tested. If all mutations were truly neutral, we would expect to call 1% or *∼* 173 of them non-neutral by chance, yielding the false discovery rate of 173*/*6971 = 2.5%.

##### 1.3.5 Growth rates of background strains in YPD

Johnson et al estimated the mean, variance and skewness of tn-insertion DFEs in 163 yeast background strains *g* in rich YPD medium, as well as the fitness *s*_*g*_ of these strains relative to a common reference strain (*39*). They also separately measured exponential growth rate *λ*_*g*_ for a subset of their background strains. Using this subset of strains, we find a very good linear relationship between *λ*_*g*_ and *s*_*g*_ in YPD (*P* = 9.632 *×* 10^−9^, *R*^2^ = 0.97),

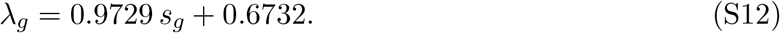

We use equation (S12) to estimate the growth rate in YPD for all 163 strains. We estimate the pivot growth rate in YPD as the growth rate at which DFE mean is predicted to cross 0 from a linear model of DFE mean against estimated background growth rate.

##### 1.3.6 Variation of growth rates and mutational effects across environments

To assess how the growth rate rank order of background strains varies across environments, we first find the median growth rate of all strains in each environment. We then call a strain as “above median” if its growth rate is above this median by at least one unit of its standard error 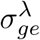 (equation (S7)). Analogously, we call a strain as “below median” if its growth rate is below the median by at least 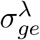. Any strain that is identified at least once as above median growth rate and at least once as below median growth rate was labelled as “Rank change” in Figure S7A).

We assessed rank-order changes of mutations across environments within each strain using an analogous procedure (Figure S8).

##### 1.3.7 Models of global epistasis

We fit equations (1) and (2) in the main text using `lm` function in R using the mean mutation effect and background strain growth rates. In addition to the estimates of slopes and intercepts, these fitted models return model *P* -values and adjusted *R*^2^ values. After Benjamini-Hochberg *P* -value correction, we find that equation (1) is significant (*P <* 0.05) for 91/94 (97%) mutations and equation (2) is significant for 90/94 (96%) mutations.

###### Accounting for measurement errors in the calculation of global epistasis slopes

As pointed out by Berger and Postma (*58*), negative slopes in equation (1) in the main text (for any given mutation *m* and environment *e*) can arise spuriously due to the fact that measurements errors in *λ*_*ge*_ and Δ*λ*_*mge*_ are correlated. We take two approaches to assess the potential impact of these correlated errors.

First, using the same approach as in Ref. (*58*), we compute the corrected correlation coefficient between Δ*λ*_*mge*_ and *λ*_*ge*_ across background genotypes for a fixed mutation *m* in a fixed environment *e* as

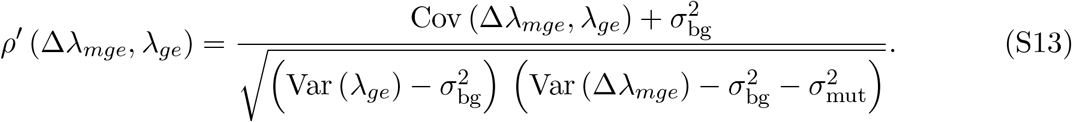

Here *λ*_*mge*_ and Δ*λ*_*mge*_ are given by equations (S6) and (S8), respectively; 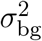 and 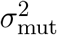 are the measurement noise variance for growth rate of the background and mutant strains, respectively, which can be calculated from expressions (S10) and (S11), respectively; and

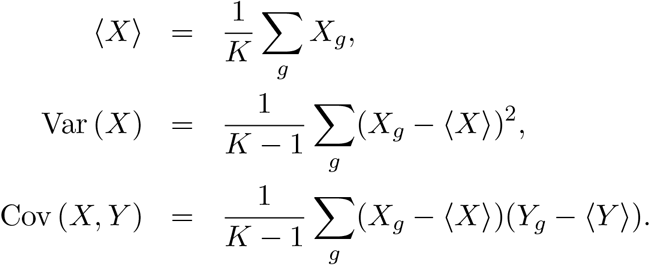

are the estimates of the mean, variance and covariance, taken over all background genotypes *g*. We find that the uncorrected correlation coefficient

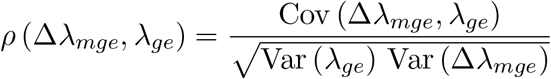

deviates very little from the corrected correlation coefficient *ρ*^*′*^ given by equation (S13) (see Figure S2D).

Second, for each mutation in each environment, we estimate the slope *b* of the regression of the growth rate *λ* of background strain against the mutant growth rate *λ*^+^ (we drop the indices for convenience). We then estimate the mutation’s global epistasis slope as *b −* 1. We find that 255/546 (47%) global epistasis slopes estimated in this way are significantly different from zero and very close to slopes estimated using equation (2) in the main text (Figure S2E).

These analyses confirm that the global epistasis trends that we observe are not spurious.

###### Comparing distribution of slopes and intercepts across environments

We tested how the distributions of fitted slopes and intercepts vary across environments using three different pairwise tests (Figure S4A).

1. A KolmogorovSmirnov test assess the overall differences between two distributions.
2. A paired t-test assess the differences between the means of the two distributions.
3. An F-test assess the difference between the variances of the two distributions.

All tests were performed using the stats package in R, and the raw *P* -values were adjusted using the Benjamini-Hochberg multiple testing correction. Adjusted *P* -values are reported in Figure S4A.

###### Comparison of the variable slopes and invariant slopes models for individual mutations

In addition to the full “variable slopes” model (equation (1) in the main text) in which a mutation can have different slopes in different environments, we also fit an “invariant slopes” model to our data in which every mutation has a single environment-invariant slope.

We compared that variable and invariant slopes models using the likelihood ratio test and found that, for the majority of mutations, we could not reject the invariant slopes model in favor of the variable slopes model which has 5 more parameters (Figure S5A). We then calculated the adjusted 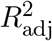 for both the invariant and variable slopes models for each mutation using the R function `lm`,

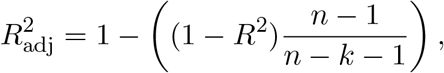

where *R*^2^ is the standard coefficient of determination, *n* is the number of observations and *k* is the number of predictors. In this case, *n* is the number of unique genotype-environment combinations in which the mutation is measured, and *k* is twice the number of environments in which the mutation is measured (variable slopes model) or the number of such environments plus one (invariant slopes model). This adjustment helps identify potential over-fitting by penalizing a high number of parameters relative to the number of observations.

To assess if the similarity of the variant and invariant slopes models could be driven by missing measurements, we apply the same procedure on the reduced data set which contains, for each mutation, only those strains for which that mutation has a measured effect in all environments. We find that the results are very similar in both data sets (Figure S5D-E).

###### Analysis of microscopic epistasis slopes

To determine whether global-epistasis slopes for a given mutation are statistically distinguishable across environments, we estimate these slopes as described above using the `lm` function in R. Along with the maximum likelihood estimates of the slopes, this function returns the standard errors of these estimates. Then, for each pairwise slope comparison, we calculate the difference between the slopes and estimate the associated error variance as the square root of the summed squared errors. Assuming that errors are normally distributed, we calculate the *P* -value for the observed error. We then apply the Benjamini-Hochberg multiple testing correction for all pairwise comparisons for a given mutation and obtain adjusted *P* -values. A pair of slopes is then called significantly different if the adjusted *P* -value is below 0.05.

To assess if the cross-environmental similarity of global epistasis slopes could be driven by missing measurements, we apply the same procedure on the reduced data set which contains, for each mutation, only those strains for which that mutation has a measured effect in all environments. We find that, similar to the full data, 87% of pairwise slope comparisons are statistically indistinguishable (Figure S5F).

We also tested whether the slope similarity was driven primarily by “flat” mutations, i.e., those where slopes were not significantly different from zero. To this end, we removed those 258/546 (47%) of mutation-environment combinations for which the slope of the fit of (1) in the main text was not significantly different from zero. Of the remaining 415 pairwise slope comparisons, 356 (86%) were not significantly different, suggesting that the “flat” mutations are not the main driver of the observed similarity of slopes across environments.

###### Determining the pivot growth rate and its variation across environments

There are two ways of determining the pivot growth rate. First, for each mutation *m* in environment *e*, we can find the background growth rate 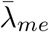 at which the mutation’s expected fitness effect is zero. Using equation (1) in the main text (with *b*_*me*_ = *b*_*m*_), we obtain an equation for 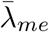,

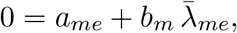

which yields 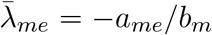. We can then obtain the pivot growth rate 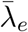 for environment *e* by averaging 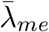 over all mutations *m*. The drawback of this approach is that it produces a value of 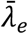 regardless of whether different mutations actually switch sign at similar background growth rates or not.

The second approach, which is the one we have taken, starts by asking whether *a*_*me*_ and *b*_*m*_ are correlated. A priori, *a*_*me*_ and *b*_*m*_ need not be correlated, or, if they are correlated, we have no a priori constrains on the slope or intercept of this correlation. In fact, we find that the relationship between *a*_*me*_ and *b*_*m*_ is very well described by the statistical model with a zero intercept and slope *c*_*e*_ *<* 0 (see Figure 2C in the main text),

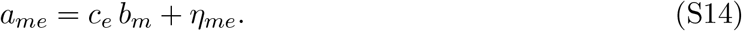

Substituting equation (S14) into equation (1) in the main text (with *b*_*me*_ = *b*_*m*_) yields

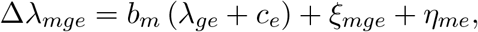

which shows that 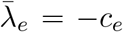, i.e., |*c*_*e*_| is the growth rate at which a typical mutation switches sign. Thus, we estimate the pivot growth rate 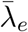 in each environment *e* as the negative of the best-fit ordinary linear regression slope in equation (S14) using the `lm` package in R.

To test if pivot growth rates significantly differ across environments, we follow a procedure analogous to the one used to test pairwise differences in microscopic epistasis slopes (see above), and we find that pivot growth rates indeed significantly differ in almost all comparisons (Figure S4B). Additionally, we find that the pivot growth rate estimated with the invariant slopes *b*_*m*_ well correlate with those estimated using variable slopes *b*_*me*_ (Figure S5C).

##### 1.3.8 Variance partitioning for the sign of mutations

Let *Y*_*mge*_ be the observed sign of mutation *m* in genetic background *g* in environment *e*, such that *Y*_*mge*_ = *±*1. The total variance in the observed signs of mutations is

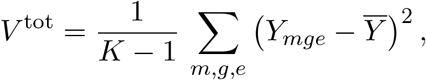

where

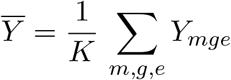

is the average sign of the mutational effect and *K* = ∑_*m,g,e*_ 1 is the total number of mutations measured across all genotypes and environments.

Let the true fitness effect of mutation *m* in genetic background *g* in environment *e* (without measurement noise) be

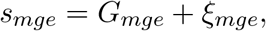

where *G*_*mge*_ = *a*_*me*_ + *b*_*me*_ *λ*_*ge*_ is the global epistasis term, *λ*_*ge*_ is the growth rate of background strain *g* in environment *e*, and *ξ*_*mge*_ is the idiosyncractic epistasis term (see equation (1) in the main text). Then, the probability that the observed sign of this mutation is positive is

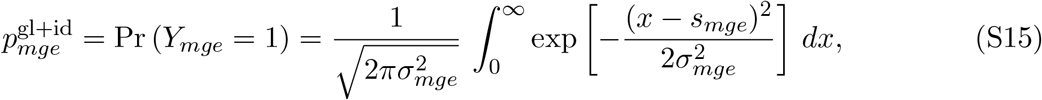

where 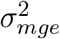 is the variance of the measurements noise for mutation *m* in background *g* in environment *e*. The super-index gl + id indicates that this probability takes into account both global and idiosyncratic epistasis. We estimate 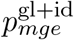 using equation (S15) with *s*_*mge*_ and *σ*_*mge*_ given by equations (S8) and (S9). This allows us to calculate the expected sign of mutation *m* in background *g* in environment *e*,

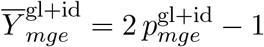

and estimate the variance in the mutational sign attributed solely to measurement noise,

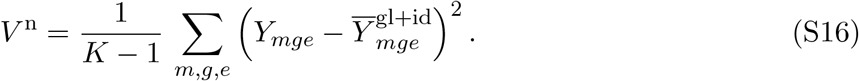

In a model without idiosyncratic epistasis, the probability 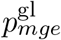 that the observed sign of the mutational effect is positive can be estimated using the same equation (S15), but with

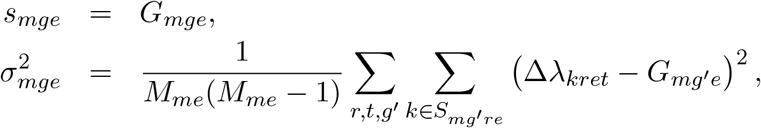

where Δ*λ*_*kret*_ are estimated with equation (S2) and *M*_*me*_ is the number of barcode-genotype-replicate-time combinations at which Δ*λ*_*kret*_ are estimated for mutation *m* and environment *e*.

Thus, the variance in the mutational signs attributed to both idiosyncratic epistasis and measurement noise is

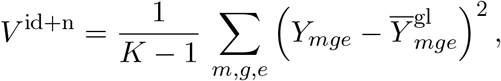

where

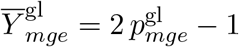

is the expected sign of mutation *m* in background *g* in environment *e* in the model without idiosyncratic epistasis. Thus, we can partition the total variance *V* ^t^ into the global, idiosyncratic and noise components as follows. Variance *V* ^n^ given by equation (S16) is attributed to noise, variance

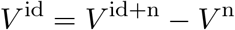

is attributed to idiosyncratic epistasis, and variance

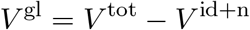

is attributed to global epistasis.

##### 1.3.9 Empirical DFEs

For YPD, we use the estimates of DFE moments and their corresponding standard errors obtained in Ref. (*39*), with minor adjustments to account for the conversion from relative fitness to absolute growth rate (see equation (S12) in Section 1.3.5). Specifically, we multiply the published DFE mean values and their standard errors by the scaling factor 0.9729 and we multiply the published DFE variance and their standard errors by 0.9729^2^. We do not adjust DFE skewness values because they are unit-less.

In every other environment *e* and for each background strain *g*, we use the fitness effect estimates Δ*λ*_*mge*_ obtained from equation (S8) to estimate the mean ⟨Δ*λ* ⟩_*ge*_, variance Var_*ge*_ [Δ*λ*] and skewness Skew_*ge*_ [Δ*λ*] of the empirical DFEs as

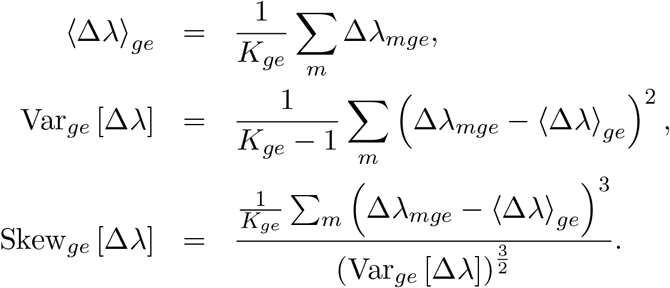

Here, *K*_*ge*_ is the number of mutations whose effects were estimated in background strain *g* in environment *e*. To estimate the uncertainty in these estimates, we resampled 70 random mutations from each empirical DFE with replacement (bootstrapping). For each resampled mutation, we drew its fitness effect from a normal distribution with mean Δ*λ*_*mge*_ (given by equation (S8)) and standard deviation 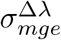 (given by equation S9). We performed 300 iterations of this procedure.

###### Sensitivity of DFE moment estimates to missing measurements

In pooled cultures, slow growing mutants may go extinct during the competition assay, which could prevent us from estimating their effects and lead to biases in our estimates of the DFE moments. In particular, highly deleterious mutations missing from the data could leads us to overestimate the DFE mean, underestimate the DFE variance and overestimate DFE skewness. Furthermore, we expect that these biases would be stronger in slower growing background strains, which could produce spurious declines in DFE mean and skewness with the background-strain growth rate. As described above, we sought to mitigate this potential issue experimentally by competing our mutants in groups with similar growth rates (see Section 1.2.3). However, these spurious effects may still be present in our data. To investigate how severe these effects might be, we plotted the number of mutations for which we have reliable fitness-effect estimates against the respective background strain growth rate. We found that the number of mutations per strain varies little with the background-strain growth rate in 30°C environments (Figure S10B), but it does vary in the expected direction in 37°C environments. However, since this analysis does not reveal the fitness effects of missing mutations, this observation alone does not imply that our estimates of DFE moments are biased for slow-growing strains in 37°C environments. Thus, we carried out three additional analyses to further probe probe the extent of these potential biases.

First, we imputed the effect of the missing mutation measurements in the 37°C environments to assess if we expect to have missed many deleterious mutations. We are only able to do this for those missing mutation-environment-strain combinations in which (i) there are sufficient measurements of the mutation in the focal environment to allow the estimation of microscopic epistasis regression, and (ii) we have an estimate of the background strain growth rate in the focal environment. Out of a total of 4252 missing mutation-environment-strain combinations, 3568 (84%) are imputable. We then calculate the predicted effect of the missing mutation-strain-environment combination in the absence of idiosyncratic epistasis using (1) in the main text with the fitted parameters and the estimated growth rate of the background strain and use it as the imputed value. We find that, while the distributions of imputed and measured fitness effects (combined across all three 37°C environments) are statistically different, the average imputed fitness effect is only by 0.002 h^*−*1^ (*P* = 7*×*10^−7^, t-test; Figure S10C), which is much smaller than 0.026 h^*−*1^, the standard deviation of the distribution of measured effects.

Second, we eliminated all strains from our analysis for which we measured less than 60 mutations (bottom 24%) and replotted DFE moments for this reduced number of strains (Figure S10D–F) and found that all the trends found in the full data set remain in this reduced data set (compare Figure S10D–F and 4 F–H in the main text).

Third, in each environment, we identified the set of 40 mutations all of which were measured in the maximum number of strains in that environment, which ranged from 16 to 40 strains, depending on environment. We then restricted our analysis of DFE moments to these strains and mutations, thereby creating a reduced data set without any missing measurements. We found that the DFE moments recapitulate the trends reported for the full data set (compare Figures S10G–I and 4 F–H in the main text).

Based on these analyses, we conclude that the trends in the DFE moments that we report are not spurious results of missing measurements.

###### Variation of the DFEs with adjusted growth rate

We carried a series of pairwise DFE comparisons across strains and/or environments. To this end, we created matched pairs of background strains using three methods:

- *Adjusted growth rate matching*. We matched each background strain *g*_1_ in environment *e*_1_ with another background strain *g*_2_ ≠ *g*_1_ in a different environment *e*_2_ */*= *e*_1_, such that the adjusted growth rate 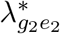 of the latter strain was most similar to the former strain among all other strain-environment combinations.
- *Raw growth rate matching*. We carried out the same procedure as above except we matched raw growth rates.
- *Strain matching*. We matched each background strain *g*_1_ in environment *e*_1_ with itself in another environment *e*_2_ such that 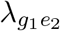 was the closes to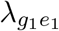.

Then, we compared the DFEs of the two matched strain-environment combinations using four metrics of similarity shown in Figure S9.

##### 1.3.10 QTL analysis

With only 42 background strains, we have little power to identify QTLs *de novo*. Instead, we assess whether certain loci identified in a previous study (*46*) help explain some of the idiosyncratic epistasis. Specifically, Bloom et al identified 37 QTLs as having significant explanatory power for the growth rate of strains in environments with pH and high temperature stress (e.g., environments YNB pH 3, YNB pH 8 and YPD 37C). Since testing the effect of all 37 loci jointly with 42 strains would lead to overfitting, we chose to restrict our analysis to the top 10 QTLs on different chromosomes^1^ that explain most variance in growth rate in any of the environments. The list of QTLs that we chose to test is provided in Table S6-Tab 2. For each QTL, we found an individual site variable in our strains that was closest to the position of the QLT peak to represent that QTL. We then tested whether these QTL-representative sites have significant explanatory power when added to our statistical model.

First, we used ANOVA to determine whether these 10 loci explain variance in background strain growth rate in each environment at a significance level of *P <* 0.05. We found that the maximum number of significant loci in any environment was two, and that the total variance in strain growth rate attributable to these QTLs ranged between 0 and 52% (Figure S11A). The strongest QTL, which explains 51.7% and 52.4% of growth rate variance in 37°C, pH 3.2 and 5.0 environments, is associated with the KRE33 locus on Chromosome 14, consistent with previous studies (*54, 39*). We then used ANOVA to determine whether the addition of these 10 loci to the generalized global epistasis model (equation (2)) significantly reduced the proportion of unexplained variance in fitness effect for each mutation. If the contribution of a locus was significant at *P* -value 0.05, then we also calculated the fraction of variance explained by this locus above and beyond the generalized global epistasis model.

Out of a total of 93 mutations in which our global epistasis models explained some variance in their fitness effects, we found three mutations (in genes UBP3 (ID 95), in GET2 (ID 61), and nearby WHI2 (ID 57)) where 27%, 23%, and 20% of overall variance was jointly explained by the 10 candidate QTLs, corresponding to 50%, 39%, and 37% of the explained variance in each mutation, respectively. For the remaining 97% (90/93) mutations, the 10 candidate loci together explain less than 11% of overall variance, with a median of only 2.1% (interquartile interval [0.8%, 3.7%]), corresponding to a median of 4.4% of the total explained variance within each mutation (interquartile interval [1.3%, 9.6%], Supplementary Table S6-Tab 9).

##### 1.3.11 GO-term enrichment analysis

In an attempt to identify possible mechanistic signals underpinning our observations, we performed a GO-term enrichment analysis. First, we tested whether our set of tn-insertions was significantly enriched for any GO terms, using the `clusterProfiler` tool and the *S. cerevisiae* reference database from R’s Bioconductor suite. We called a GO term as enriched if it passed the 0.05 significance threshold after the Benjamini-Hochberg multiple testing correction. We found 11 significantly enriched GO terms, including regulation of transcription by RNA polymerase II and double-strand break repair via nonhomologous end joining, with the enrichment ranging from 3-to 10-fold compared to the random expectation (Table S4). Thus, our set of mutations is not entirely random, which is expected given that we have excluded lethal and unconditionally neutral mutations (see Section 1.1.2).

We then asked whether mutations with certain global microscopic epistasis slopes were enriched for some GO terms relative to the overall set. To this end, we separated mutations into into four groups, based on the quantiles of the distribution microscopic epistasis slopes *b*_*m*_ shown in Figure 2A in the main text. Then, for each group, we generated the empirical null distribution of each GO-term count by randomly resampling GO terms 10,000 times from the set of terms for all mutations. This allowed us to calculate the empirical *P* -value for all observed GO-term counts in each group. After applying the Benjamini-Hochberg multiple testing correction, we found no terms in any group that were either significantly enriched or depleted relative to the full set of mutations.

#### 1.4 Theoretical calculation of DFE moments

Here we derive the moments of the distribution of fitness effects (DFE) of mutations from the generalized global epistasis equation (equation (2) in the main text). Since we consider the environment fixed, we drop the subindex *e*. To simplify notations, we will denote the adjusted growth rate of the background strain *g* by 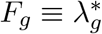 and we denote the fitness effect of a mutation in this background by *s*_*g*_ *≡* Δ*λ*_*g*_. To derive DFE moments, we assume that the effects of mutations *s* are drawn from a continuous distribution defined by equation

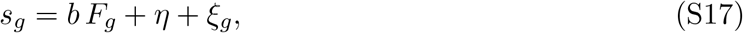

which is the continuous analog of equation (2) in the main text. Here *b* and *η* are the slope and the *y*-intercept of the focal mutation, and *ξ*_*g*_ is the idiosyncratic epistasis of this mutation in the background strain *g*. We assume that *b* and *η* are independent, and that *b* has probability density *p*_sl_, *η* is normally distributed with zero mean and variance 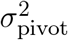. We also assume that *ξ*_*g*_ is normally distributed with zero mean and variance 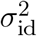 (which can in principle depend on *b*, see Ref. (*44*)). Thus, conditional on *b* and *η*, the fitness effects *s*_*g*_ of the mutation in the background *g* is a normal random variable with distribution with mean ⟨*s*|*b, η*⟩ = *b F*_*g*_ + *η* and variance 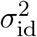. Then, the DFE *p*_*g*_(*s*) in the background *g* with adjusted growth rate *F*_*g*_ is given by

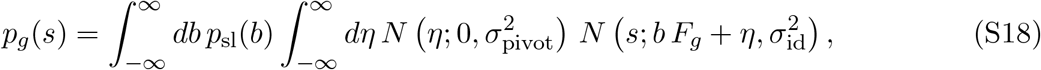

where *N* (*x*; *µ, σ*^2^) is the normal probability density with mean *m* and variance *σ*^2^. Since

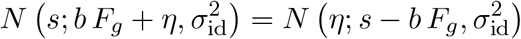

and since

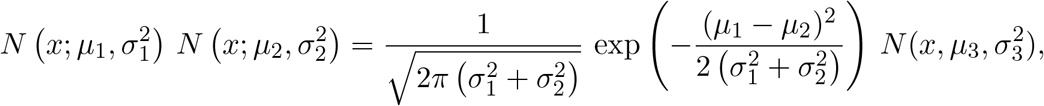

where 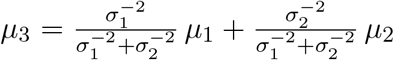 and 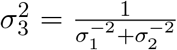 for any *µ*_*1*_, *µ*_*2*_, 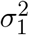 and 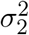, the integral with respect to *η* can be taken, such that the expression (S18) simplifies to

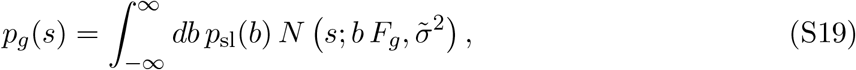

where we denoted 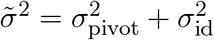.

Equation (S19) allows us to compute the DFE mean as

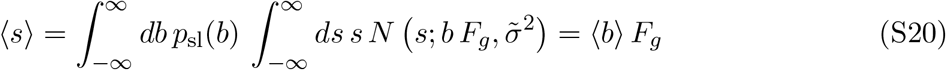

and higher central moments of the DFE as

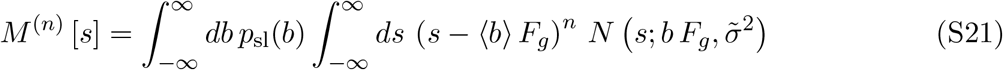

Using expression (S21) and the fact that

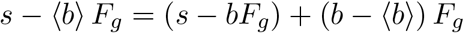

we obtain the following explicit expressions for the DFE variance Var [*s*] and its third central moment *M* ^(3)^ [*s*],

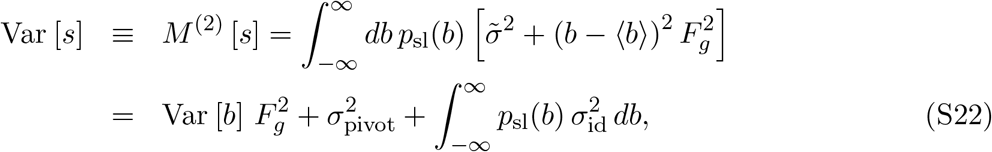

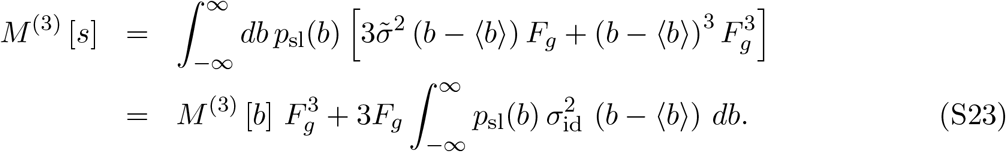

Here Var [*b*] and *M*^(3)^ [*b*] are the variance and the third central moment of the distribution *p*_sl_(*b*) of global epistasis slopes, respectively.

We find that the variance of residuals and slopes are correlated (Figure S4D). Thus, we set 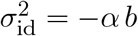, and the expressions (S22) and (S23) become

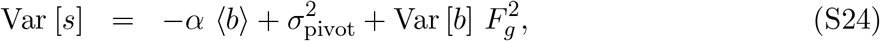

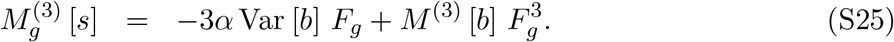

and the skewness of the DFE is given by

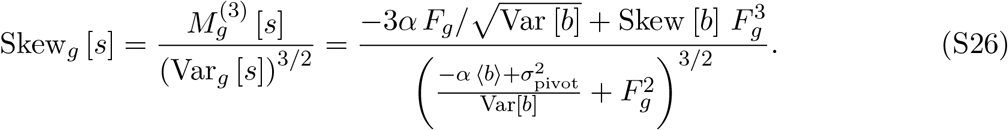

Equations (S20), (S24) and (S26) show that when *F*_*g*_ = 0, DFE mean and skewness are zero and the DFE variance achieves its minimal value 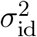. Furthermore, since Skew [*b*] *<* 0, DFE skewness monotonically declines from *−*Skew [*b*] *>* 0 to Skew [*b*] *<* 0.

To show that all odd moments of the DFE vanish when *F*_*g*_ = 0, we notice that, when *F*_*g*_ = 0, equation (S19) simplifies to

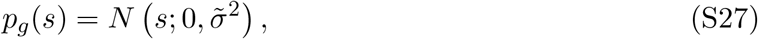

i.e., the DFE is a normal distribution, which implies that all its odd central moments vanish. To show that all even moments of the DFE reach their minimum at *F*_*g*_ = 0, we differentiate expression (S21) with respect to *F*_*g*_ at *F*_*g*_ = 0 and obtain

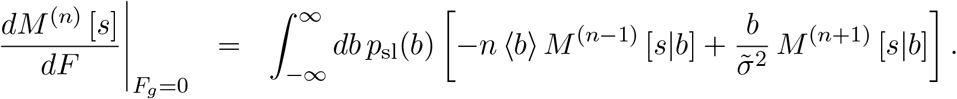

where 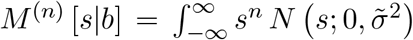 *ds* is the *n*th central moment of a normal distribution with mean zero and variance 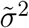 Therefore, when *n* is even,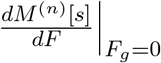 vanishes because all odd central moments of a normal distribution are zero. To see that *M* ^(*n*)^ [*s*] achieve their minimum at *F*_*g*_ = 0 for any even *n*, we find that the second derivative of *M* ^(*n*)^ [*s*] at *F*_*g*_ = 0 is given by

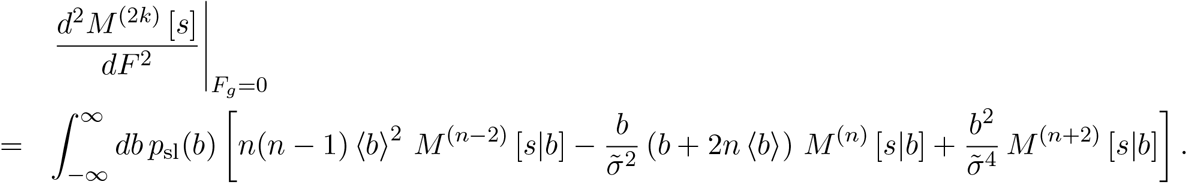

The even moments *M* ^(*n*)^ [*s*|*b*] of the normal distribution 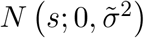 can be expressed as 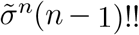 where (*n −* 1)!! = (*n −* 1)(*n −* 3) *· · ·* 3 *·* 1 is the double factorial. Therefore, we have

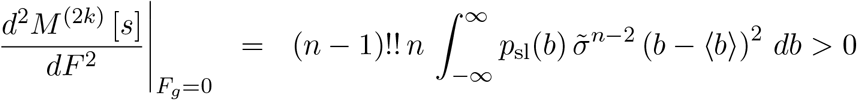

for any even *n*, which implies that *M*^(2*k*)^ [*s*] indeed achieves its minimum at *F*_*g*_ = 0.

##### Parameterization of DFE moments

The predicted DFE moment curves in Figure 4C–H are calculated with equations (S20), (S24), and (S26) for DFE mean, variance and skewness, respectively. These equations depend on the moments of the distribution of slopes *p*_sl_(*b*) and parameter *α*, which determines how variance in idiosyncratic epistasis depends on the global-epistasis slope. We estimated the moments of *p*_sl_(*b*) from the empirical distribution of global epistasis slopes obtained from fitting equation (2) in the main text (see Figure 2A). We estimated the value of the parameter *α* = 1.37*×*10^−3^ by regressing the variance 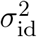 of the idiosyncratic epistasis residuals *ξ*_*mge*_ against the microscopic epistasis slopes *b*_*m*_, with the regression constrained to pass through the origin (see Figure S4D).

To quantify how well these theoretical predictions fit our data, we compare them to the best-fit polynomial of the corresponding degree (Figure 4F–H). Specifically, for mean and variance, we use the `lm` function in R to fit a first and second degree polynomial, respectively. For skewness, we utilize the predicted variance values from the fit of the second degree polynomial to parameterize the denominator of the rational function in equation (S26). We then fit the best fit third degree polynomial in the numerator. The fitted and theoretically predicted parameters are shown in Table S3.

### 2 Supplementary Text

#### 2.1 Justification for using absolute growth rate rather than relative fitness

As far as we know, almost all previously reported instances of global epistasis were observed with respect to competitive fitness in batch culture (but see Ref. (*19*)). In contrast, we designed our study so that we can estimate not only competitive fitness but also the absolute growth rates of our background strains and mutants. In some ways, these two choices are equivalent. Specifically, (i) the fitness effects of mutations given by equation (S8) in fact depend only on the relative barcode frequencies (see equation (S2)); and (ii) the competitive fitness of background strains relative to each other can be calculated by taking differences between their estimated absolute growth rates. Thus, most of our results would not be impacted by our choice. However, we see three advantages of using the absolute growth rate.

Using absolute growth rate makes our findings independent of any particular reference strain and therefore potentially more generalizable to other systems.

To the best of our knowledge, almost all previous observations of global epistasis were made with respect to competitive fitness. However, competitive fitness in batch-culture conditions depends (non-linearly) on multiple components, such as the duration of the lag phase, exponential growth rate, survival during starvation, etc. (*59, 60, 61, 62*). Thus, it is conceivable that epistasis could arise at the level of competitive fitness even among mutations that exhibit no epistasis for any of the individual components of fitness. Our observation that global epistasis holds at the level of growth rate strongly suggests that it is an intrinsic property of the cell. This narrows the scope of possible mechanistic explanations for this phenomenon and suggests that it would be relevant in any environment where growth is an important component of fitness.

Absolute growth rate provides us with a concrete threshold for strain viability. In other words, there is no a priori minimum on a relative growth rate scale, whereas the absolute scale naturally cuts off at zero. This fact allows us to attribute meaning to the intercepts of microscopic global epistasis regressions and to the location of the pivot growth rate relative to zero. In particular, a model in which global epistasis lines intersect below viability would have been consistent with our a priori intuition that most mutations are unconditionally deleterious and that DFEs for all viable genotypes are negatively skewed.

#### 2.2 Comparison with previous studies and potential limitations

As mentioned in the main text, many studies carried out in several microbial species found evidence of microscopic global epistasis (*11, 12, 34, 35, 36, 37, 38, 39, 40, 41, 42*). There are also a few recent studies that found evidence for macroscopic global epistasis (*31, 33, 34, 39*). It is instructive to compare our results with these previous works. We focus here on the four most relevant studies, Refs. (*19, 31, 33, 34*), grouped by the types of mutations that they have measured.

##### Diaz-Colunga et al, Ref. (*19*)

Diaz-Colunga et al examined how the patterns of microscopic global epistasis within a combinatorially complete set of mutations in the *DHFR* gene (from *Plasmodium falciparum*, integrated into the genome of *S. cerevisiae*) vary across environments where the concentration of the antimalarial drug pyrimethamine varies over five orders of magnitude. They found that the patterns of global epistasis vary across this wide range of drug concentrations. However, interestingly, these patterns remain largely invariant for concentrations below 1 µM, despite the fact that the growth rates of mutants and their rank order change over this range (*63*). There are many differences between our experimental design and that of Ref. (*19*), e.g., types and number of mutations, background strains, type and degree of environmental variation. So, it is difficult to draw definitive conclusions. However, one intriguing possibility is that the near-invariance of the global epistasis patterns across environments has to do with how distributed or localized the environmental perturbation is. For example, it is possible that at low concentrations (below 1 µM) the drug exhibits a global toxic effect on the cell, similar to our temperature and pH perturbations, and global epistasis patterns remain largely invariant. At high concentrations (above 1 µM), the toxicity may come primarily from the drug’s interaction with its target protein, dihydrofolate reductase enzyme, and the patterns of epistasis become highly idiosyncratic.

##### Aggeli et al, Ref. (*31*)

Aggeli et al used the barcode-lineage tracking approach (*29*) to measure how the beneficial part of the DFE (bDFE) varies across four strains of *S. cerevisiae*, one ancestral strain and three adapted mutants each carrying a single beneficial mutation. They found that the bDFE in all three adapted strains is shifted left relative to that of the ancestral strain, which is qualitatively consistent with our results.

##### Johnson and Desai, Ref. (*34*), and Couce et al, Ref. (*33*)

Both Johnson and Desai and Couce et al use the transposon-insertion mutagenesis followed by sequencing (TnSeq) approach to measure how the DFE changes along an evolutionary trajectory over about 10 to 15 thousand generations. While conceptually very similar, these studies differ in two aspects. First, Johnson and Desai sampled strains from six time points of the long-term evolution experiment carried out in budding yeast *S. cerevisiae* in two environments, YPD 30°C and SC 37°C^2^, whereas Couce et al sampled strains from two to three time points of the Lenski’s long-term evolution experiment (LTEE) in bacterium *Escherichia coli*. The second difference is that Johnson and Desai used the same TnSeq approach as we employed in the present study, i.e., they measured the effects of the same *∼* 100 tn-insertions, whereas Couce et al used “saturated” TnSeq approach, with *∼* 10^5^ tn-insertions covering 78% of all *E. coli* genes.

Differences in experimental design constrained the power of the two studies in different ways. While both studies found the vast majority of tn-insertions to be deleterious, Couce et al were able make statistical statements about beneficial mutations whereas Johnson and Desai focused only deleterious mutations. Johnson and Desai found evidence for global microscopic epistasis in both environments, although the fraction of variation explained by background fitness was generally smaller than in Ref. (*39*) where the background strains were derived from the RM*×*BY cross, as in the present work. Couce et al found significant differences in the effects of many tn-insertions across genetic backgrounds (i.e., evidence of microscopic epistasis), but they likely did not have enough power to detect any global component in it. However, microscopic global epistasis has been previously observed among adaptive mutations in the LTEE (*11, 43*).

Focusing now on macroscopic epistasis, Couce et al found that evolved clones had a clearly depleted bDFE compared to the LTEE ancestor, which is consistent with our model as well as with previous observations of “the rule of declining adaptability” (*64, 65*). However, they detected no systematic shifts in the overall DFE over the course of evolution, which is inconsistent with our results. Johnson and Desai found fitness-dependent shifts in the DFE among clones evolved in the YPD 30°C consistent with our results and Ref. (*39*). However, despite the presence of global microscopic epistasis, they found no consistent fitness-dependent shifts in the DFE among clones evolved in SC 37°C, similar to the results by Couce et al.

One possible technical reason that could explain the lack of a detectable shift in the DFE in the Couce et al study is that their DFEs appear to be dominated by neutral or nearly neutral mutations, similar to the “large library” investigated in Ref. (*39*). If the majority of mutations are in fact unconditionally neutral (or at least neutral in all investigated strains), detecting shifts in the distribution of fitness effects of non-neutral mutations becomes statistically challenging. However, this cannot explain the why Johnson and Desai failed to observe DFE shifts in the SC 37°C environment because they specifically focused on non-neutral mutations and had power to detect such shifts.

Patterns of global epistasis could be fundamentally different between *S. cerevisiae* and *E. coli* or they might differ between exponential growth (used in our study) and batch culture fitness (used in both Couce et al and Johnson et al). Yet, neither of these explain the puzzling differences between yeast strains evolved in two environments observed by Johnson and Desai. Perhaps a more likely biological explanation for the aforementioned discrepancies is that epistasis involving mutations sampled along an evolutionary trajectory may be different than epistasis involving random genetic variants (*66*). The background strains used in this study are hybrids derived from the RM*×*BY cross and may represent more or less random genetic variation^3^, or a least variation that has not been under selection in our experimental conditions. In contrast, background genotypes in the studies by Johnson and Desai and by Couce et al were sampled from adapting populations and thus represent selected (i.e., non-random) genetic variation. Moreover, mutations that drive adaptation to different environments may have different physiological effects and, as a result, exhibit different types of epistasis. Indeed, Johnson and Desai speculate that adaptation to YPD 30°C may be primarily driven by mutations in core cellular processes related to growth while adaptation to SC 37°C may be driven at least in part by mutations improving survival during heat stress; they then argue that these types of mutations could plausibly exhibit different types epistasis, which could explain the observed discrepancy between conditions.

Another intriguing possibility is that evolved genotypes could have experienced second-order selection for higher evolvability and/or lower robustness. In fact, such selection has been documented in the LTEE (*67*). Generally, second-order selection favors strains with DFEs that have heavier beneficial tails and/or weaker deleterious tails. Thus, even if the overall fitness landscape is characterized by the statistical trends shown in Figure 4 in the main text, second-order selection could in principle favor genotypes that deviate from these trends.^4^ Of course, the degree of such bias would be determined by the amount of DFE variation available in the population and on the strength of second-order selection, both of which could vary between environments. This could explain difference between the results of our study and those of Couce et al and Johnson and Desai.

Overall, this discussion highlights that extrapolating our results to predict the outcomes of evolution may not be straightforward. We interpret our results as revealing the overall statistical structure of genotype-to-fitness maps (see Section 2.3). How evolution navigates populations through these maps, especially in the presence of second-order selection or environmental variability, is not yet understood. These questions represent an interesting area for future research.

##### 2.3 Connection between the generalized global epistasis equation and deterministic genotype-to-fitness maps

Following Reddy and Desai (*44*) we interpret the generalized global epistasis equation (2) in the main text as a statistical description of an underlying deterministic genotype-to-fitness map (GFM), also called a fitness landscape. Considering a fixed environment, we assume that every genotype *g* has a particular growth rate *λ*_*g*_. Mathematically, this means that there exists a function *F* : *G →* ℝ^+^ from the genotype space *G* to the positive semi-axis ℝ^+^ representing growth rate. We denote the space of all GFMs (for a given genotype space *G*) as *ℱ*. Reddy and Desai show that there exists a subset of GFMs *ℱ*_GE_ *⊂ ℱ* in which the fitness effects of almost all mutations decline with the fitness of the background genotype (*44*) as in Figure 4A in the main text. In this section, we first demonstrate the existence of such GFMs for a simple 4-site genome and provide an intuition for how to interpret equation (2) in the main text. Then, we discuss the connectedness model developed by Reddy and Desai that allows us to construct a GFM for a genome of any length where patterns of global epistasis quantitatively match those observed in the data.

###### 2.3.1 Four-site GFM with global epistasis

Figure S12A shows a GFM for a four-site genome where each site (or locus, or position) carries one of two alleles, denoted by *−*1 and +1. To make a connection with our tn-mutations, we can imagine each position to be a gene and interpret the alleles +1 and *−*1 as the presence and absence of a functional gene at that position, respectively^5^. There are a total of 16 genotypes that we refer to by letters *a, b, c*, …, *p* (Figure S12A), and each genotype is assigned a fitness value. We consider only those mutations that change the allele at a single locus (e.g., a change from *−*1 to +1 at position 3), such that each mutation can be interpreted as either gene gain (a change from *−*1 to +1) or gene loss (a change from +1 to *−*1). The fitness effect of a mutation is simply the difference in fitness of the two genotypes connected by that mutation. This GFM has two fitness peaks (i.e., genotypes in which all mutations are deleterious), a local fitness peak at genotype *d* (*λ* = 0.623) and the global peak at genotype *o* (*λ* = 0.65).

Figure S12B shows that both gene-gain and gene-loss mutations exhibit global epistasis patterns with negative slopes on this GFM. We can now use this GFM to gain an intuition for how to interpret these global epistasis patterns.

Since the fitness effect of a mutation is simply a difference of the corresponding fitness values, the fitness effect of an immediate reversion of a mutation is of course the negative of the fitness effect of the mutation itself. For example, the loss of gene 1 in genotype *c* = (1, 1, *−*1, 1) provides a fitness benefit of +0.411 and results in genotype *j* = (*−*1, 1, *−*1, 1); immediately reverting that mutation (i.e., gaining gene 1 back) naturally has a fitness cost of *−*0.411 (see labelled mutations in Figure S12B). However, if the reversion is not immediate, i.e., if it occurs after one or more intervening mutations, then, due to epistasis, its effect will in general no longer be equal to the negative of the effect of the original mutation. For example, if genotype *j* gains gene 3 before regaining gene 1, the gain of gene 1 becomes beneficial with the fitness effect +0.075 (because it now occurs in the new genetic background *e* = (*−*1, 1, 1, 1)).

Based on the relationship between the fitness effects of mutations and their immediate reversions, one might naively expect that gene losses and gene gains would exhibit patterns of global epistasis with slopes of the opposite sign. However, as Figure S12B demonstrates, this need not be the case. In fact, how the global epistasis slope of a given mutation relates to the slope of its reversion is in general unclear. But since the two alleles *±*1 are simply arbitrary labels and therefore exchangeable, the statistical distributions of slopes of the two mutation types must be identical.

The fact that gene gains and losses are equivalent in this model leads to another somewhat counter-intuitive observation, namely that both gene gains and losses are beneficial for low-fitness genotypes and deleterious for high-fitness genotypes. For example, consider two genotypes with almost identical low fitness of about 0.17: genotype *c* that possesses gene 1 and genotype *h* = (*−*1, 1, 1, *−*1) that lacks it (Figure S12A). As mentioned above, a loss of gene 1 would provide genotype *c* with a fitness advantage of +0.411. At the same time, gaining gene 1 would also provide genotype *h* with a fitness gain of +0.137 (Figure S12B). Thus, whether a gain or a loss of a given gene is beneficial depends on the presence and absence of other genes. In other words, within this model, it is impossible to ascertain whether having certain genes is always “better” than not having them.

Finally, one can use the global epistasis lines in Figure S12B to make general predictions about adaptive evolution. Consider, for example, genotype *c* = (1, 1, *−*1, 1) with *λ* = 0.172 as the starting genotype. It has access to four mutations, gene losses at positions 1, 2 and 4 and gene gain at position 3. Loss of gene 4 is expected to be unconditionally deleterious (the dashed orange line is below zero). Since genotype *c* is below the pivot growth rate for all other mutations and since both loss of gene 1 and gain of gene 3 have the steepest slopes, we would expect that these two mutations would be the most beneficial, providing expected fitness gains of about +0.3 and +0.2, respectively. Thus, we would expect that one of these mutations would fix in the first adaptive step. Once it does, the other mutation is expected to become close to neutral; the only remaining strongly beneficial mutation would then be the loss of gene 2 that would provide a fitness gain of about 0.15. Once gene 2 is lost, we would expect adaptation to stop at a final fitness value between about 0.50 and about 0.62 and resulting either in a genotype with one gene (gene 4) present or in a genotype with one gene (gene 2) absent. These predictions are broadly borne out by the examination of the full GFM depicted in Figure S12A.

##### 2.3.2 A deterministic fitness landscape that recapitulates empirical patterns of global epistasis

As mentioned above, the connectedness (CN) model developed by Reddy and Desai (*44*) allows us to sample GFMs that exhibit empirically observed patterns of global epistasis. Before describing the properties of this model, it is important to make two general remarks. First, it is clear that not all GFMs exhibit global epistasis, i.e., *F*_GE_ is a strict subset of *F*. For example, on additive fitness landscapes, the fitness effect of each mutation is constant across genetic backgrounds, implying that such GFMs do not belong to ℱ _GE_. Thus, constructing GFMs that exhibit global epistasis is not entirely trivial. However, the CN model as well as related models (*68*) accomplish this. Let us denote the space of GFMs produced by the CN model as ℱ _CN_. Thus, ℱ _CN_ *⊂* ℱ _GE_.^6^ Second, the space of GFMs ℱ _GE_ that exhibit global epistasis is not fully characterized. For example, it is unclear how big it is or how well the GFMs produced by the connectedness model cover it. It is entirely possible that ℱ _CN_ is a highly biased subset of ℱ _GE_, such that, for example, GFMs produced by an empirically parameterized CN model may not contain the true GFM that generated the data.

###### Description of the connectedness (CN) model

In this model, an organism’s genotype is represented by a series of diallelic loci with alleles denoted as +1 or *−*1. Each locus *i* is parameterized by a single parameter *µ*_*i*_ which is the probability for that locus to participate in each of *M* “pathways”. Thus, in each realization of this model, each locus *i* is randomly assigned to a number of pathways. Each pathway *p* is described by a quantitative phenotype *y*_*p*_ whose value is determined by equation (2) in Ref. (*44*),

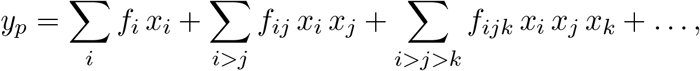

which depends on the alleles *x*_*i*_ = *±*1 at all participating loci. In other words, all loci that belong to the same pathway interact with each other through pairwise, triple, quadruple, etc. interactions. In each realization of the model, the interaction coefficients *f*_*i*_, *f*_*ij*_, *f*_*ijk*_, … for each pathway *p* are drawn randomly from the standard normal distribution. The fitness of the organism is then given by the sum of *y*_*p*_ across all pathways. Thus, each realization of the CN model produces a canonical GFM, akin to the one depicted in Figure S12A.

###### Fitting the CN model to data

Reddy and Desai show that when the number of loci is large, the global epistasis slope for a “directionless” mutation at locus *i* (i.e., either a flip from *−*1 to 1 or vice versa) is given by

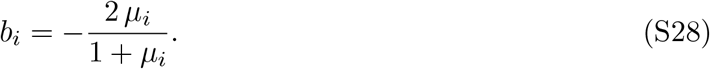

To fit the *µ*_*i*_ parameters to our data, we consider a genome with 94 loci, corresponding to the tn-insertion locations. Since this model considers directionless mutations, we follow Reddy and Desai and first calculate the global epistasis slopes in the concatenated data set which contains both the effects of tn-insertions and the corresponding effects of tn-reversions (*44*). The distribution of slopes in concatenated data in 30°C pH 5.0 is shown in Figure S13A and the corresponding ‘bowtie’ plot is shown in Figure S13D. We then use equation (S28) to fit the parameter *µ*_*i*_ for each locus using these empirical slopes. We then generate a particular instance of the CN-GFM using the code provided with Ref. (*44*), where we set *M* = 200. In this computational implementation, if a pathway is populated by less than one or more than 14 loci, all pathway members are resampled. After this assignment, an average pathway contains approximately 11 loci. This code also normalizes the fitness values so that the total variance in fitness equals one. Because the fitness values produced by this model have no biological meaning, we then linearly shift and scale them to match our data. Specifically, we set the mean fitness of all genotypes to 0.245 which is the pivot growth rate in our 37°C pH 5.0 environment (Figure S13E). We also scale all fitness values by factor 0.2, which defines the variance in fitness and thereby allows us to match the empirical slope-intercept correlation (Figure S13B–D).

###### Correlation length in the empirically parameterized CN model

Having obtained an empirically parameterized GFM, we next sought to understand its properties. While a thorough investigation of this fitness landscape goes beyond the scope of this work, one property of interest is the degree of ruggedness, which can be measured by the correlation *γ*_*d*_ between the fitness effects of mutations introduced into two genotypes separated by *d* mutational steps (*69*). For any GFM, *γ*_0_ = 1 by construction and how it changes with *d* is indicative of the degree of landscape ruggedness. At one extreme, for an additive landscape, *γ*_*d*_ = 1 independently of *d*. At the other extreme, for a House-of-Cards fitness landscape (also known as the “uncorrelated” fitness landscape (*70*)), where the fitness of every genotype is assigned independently, *γ*_*d*_ = 0 for any *d ≥* 1.

To calculate *γ*_*d*_ for a given initial genotype *g*_0_, we first retrieve the effect of all single mutations in that background genotype. Then, for each locus, we calculate the effect of mutating this locus in each of the remaining 93 single-mutant neighbors of *g*_0_. *γ*_1_ is the correlation coefficient between the fitness effect of mutating a locus in the original background genotype *g*_0_ and in a single-mutant neighbor of *g*_0_. We calculate *γ*_*d*_ for *d* = 2, 3, …, 20 in the same way, except now, for each focal locus *i* whose effect is being assessed, we calculate the effect of *i* in 93 background genotypes that are sampled from the *d*-neighborhood of *g*_0_. These *d*-neighbors are sampled while the allele at locus *i* is kept fixed at its *g*_0_ value. To ensure that the observed correlation structure is consistent across different locations in the landscape, we calculate *γ*_*d*_ for three initial background genotypes *g*_0_, whose initial fitness were chosen to be low (*λ* = 0.02, yellow point in Figure S13E), intermediate (*λ* = 0.24, cyan point in Figure S13E) and high (*λ* = 0.63, red point in Figure S13E).

We find that the fitness-effect correlation *γ*_*d*_ decays on our empirical GFM roughly exponentially, with decay rate of about 0.15, i.e., halving approximately every 4.6 mutational steps (Figure S13F). This correlation length is much longer than for the House-of-Cards model, but it is much shorter than the genetic distances of *∼* 10^4^ SNPs that separate background strains in our data set. The latter is not surprising since we parameterized the CN model using only the 94 tn-insertion loci. Thus, this estimate of correlation length may not capture the scale of correlations on the true yeast GFM (for example, in the real yeast genome there could be many neutral non-epistatic mutations that greatly extend the correlation length); rather, it simply illustrates that an GFM consistent with our data displays an intermediate degree of ruggedness.

###### Evolutionary simulations on an empirically parameterized CN model

The generalized global epistasis equation alone allows us to simulate evolution at the fitness level. For example, several previous studies have shown that global epistasis is sufficient to recapitulate fitness trajectories observed in evolution experiments (*20, 25, 44*). However, reconciling fitness trajectories with evolutionary dynamics observed at the genetic level is harder (*20*). To understand how evolution unfolds at the genetic level, we need an explicit GFM model, such as the one generated by the CN model described above.

If we interpret alleles +1 and *−*1 as the presence and absence of functional genes, respectively (so that a mutation from +1 to *−*1 corresponds to a gene loss and a mutation from *−*1 to +1 corresponds to a gene gain^5^), we can begin asking interesting questions about the evolution of genome composition in a population or, more generally, the evolution of the pan-genome at the level of a species (*71*). One simple question we can ask is how many (non-essential) functional genes an organism will retain if the fitness effects of these mutations exhibit global epistasis.

To explore this question, we ran 50 replicate Wright-Fisher simulations with population size *N* = 10^4^ and mutation rate *U* = 10^−5^ in the strong selection weak mutation regime, starting either with a random low-fitness genotype (*λ* = 0.02, with 53/94 initially intact genes) or with a genotype with all genes intact (i.e., a genotype with with +1 alleles at all loci), which has initial fitness of *λ* = 0.18. As expected, populations starting with both initial genotypes gain fitness at a characteristically decelerating pace (Figure S13G,I). However, they approach different sets of fitness peaks, with populations starting with all intact genes on average approaching a plateau at fitness of about 1.40 (Figure S13G) and those starting with the low-fitness genotype on average approaching a plateau at fitness of about 1.50 (Figure S13I). The differences between the genome dynamics of these two ensembles are even starker (Figures S13H,J). In populations that start with the low-fitness genotype, the number of disrupted genes quickly plateaus at about 43/94 (46%) (Figure S13J), whereas populations that start with all intact genes, the number of gene disruptions continues to accumulate over thousands of generations and reaching on average approximately 17/94 (18%) after 50, 000 generations and appearing to plateau near 20/94 (21%, Figure S13H).

The fact that the evolutionary trajectories starting at different initial genotypes approach different sets of fitness peaks confirms our prior observation (Figure S13F) that the CN-generated fitness landscape is quite rugged. In fact, statistical differences at the fitness and genetic levels between evolutionary trajectories that originate at different initial genotypes could be used to experimentally test how accurately the CN-generated GFM captures the true underlying GFM. Our simple simulations also suggest that an empirically parameterized CN model can potentially be used to explain properties of pan-genomes, such as the existence and the size of the accessory genome, the degree variation in genome content among populations, etc. (*71*).

##### 2.3.3 Does the Reddy-Desai theory accurately describe biological organisms?

All the results presented in this paper are remarkably consistent with the Reddy-Desai theory (*44*). Does this consistency imply that this theory in fact provides a good description of biological organisms, at least for understanding how mutations affect fitness? As far as we are aware, this theory (along with the related model by Lyons et al (*68*)) is the only one available for explaining the observed global epistasis patterns. This theory is general, self-consistent and does not appear to lead to any pathological or paradoxical behaviors, as we have shown above in this section. However, one conceptual problem is that the Reddy-Desai theory is based on mapping genotypes to fitness directly, without accounting for organismal physiology that underlies fitness. Global epistasis (with negative slopes) arises in this theory as a consequence of the central limit theorem when sites are involved in many uncorrelated genetic interactions. As such, global epistasis in this model is in essence regression to the mean. But in real biological organisms, sites or genes do not interact randomly but rather because they are involved in physiological processes that map lower-level functions (e.g., enzyme activities) to higher-level functions (e.g., metabolic fluxes) in a non-linear way (*43, 51*). It is a priori unclear if and under what conditions would such physiologically-grounded genetic interactions be uncorrelated. Thus, we see it as an important open problem to propose physiologically grounded models that are consistent with the Reddy-Desai theory or alternatively reproduce global epistasis via mechanisms other than regression to the mean.

## 3 Supplementary figures

**Figure S1.**
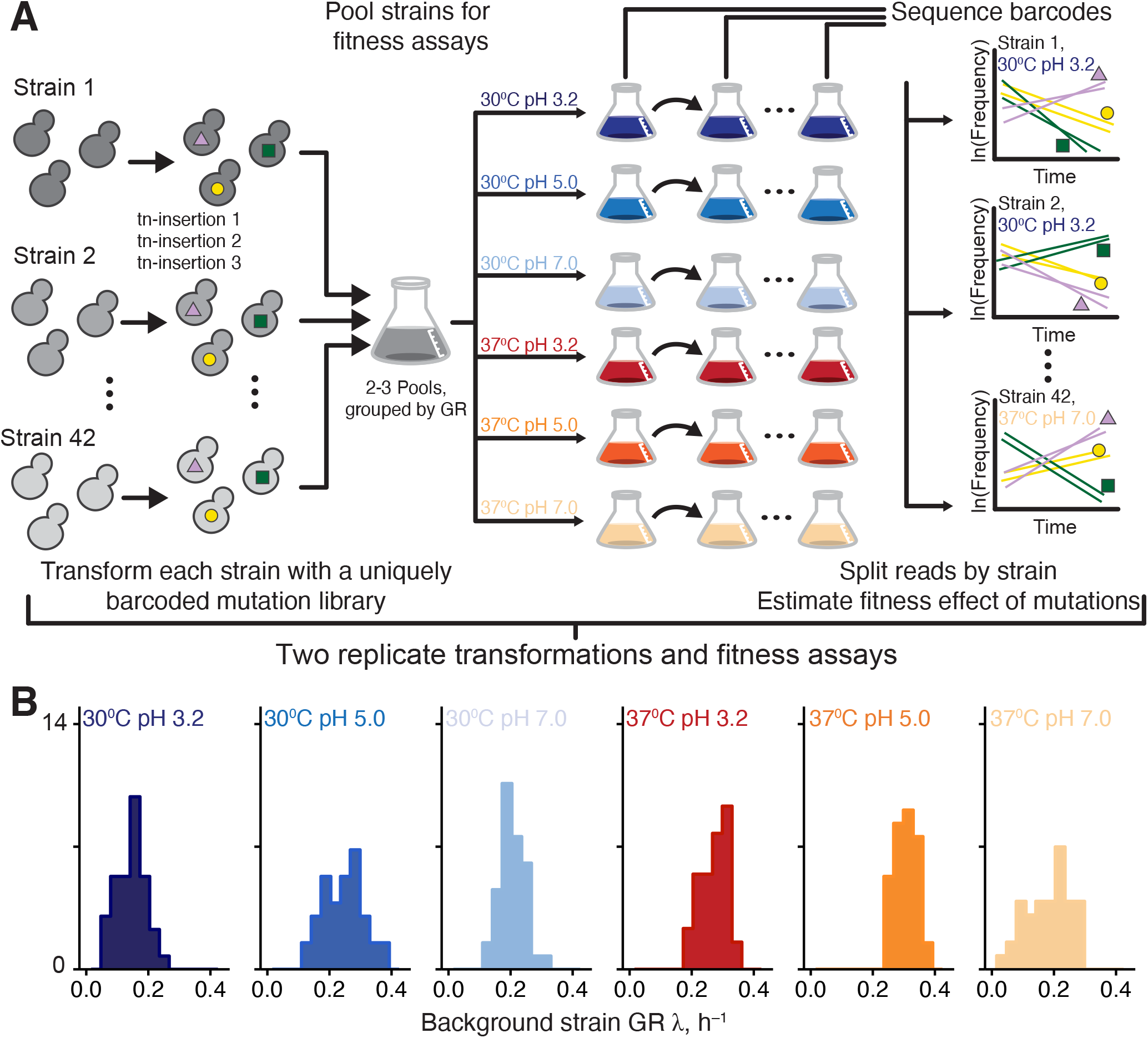
Experimental setup. **A**. Schematic of the experiment. **B**. Distribution of growth rates of background strains in all environments.

**Figure S2.**
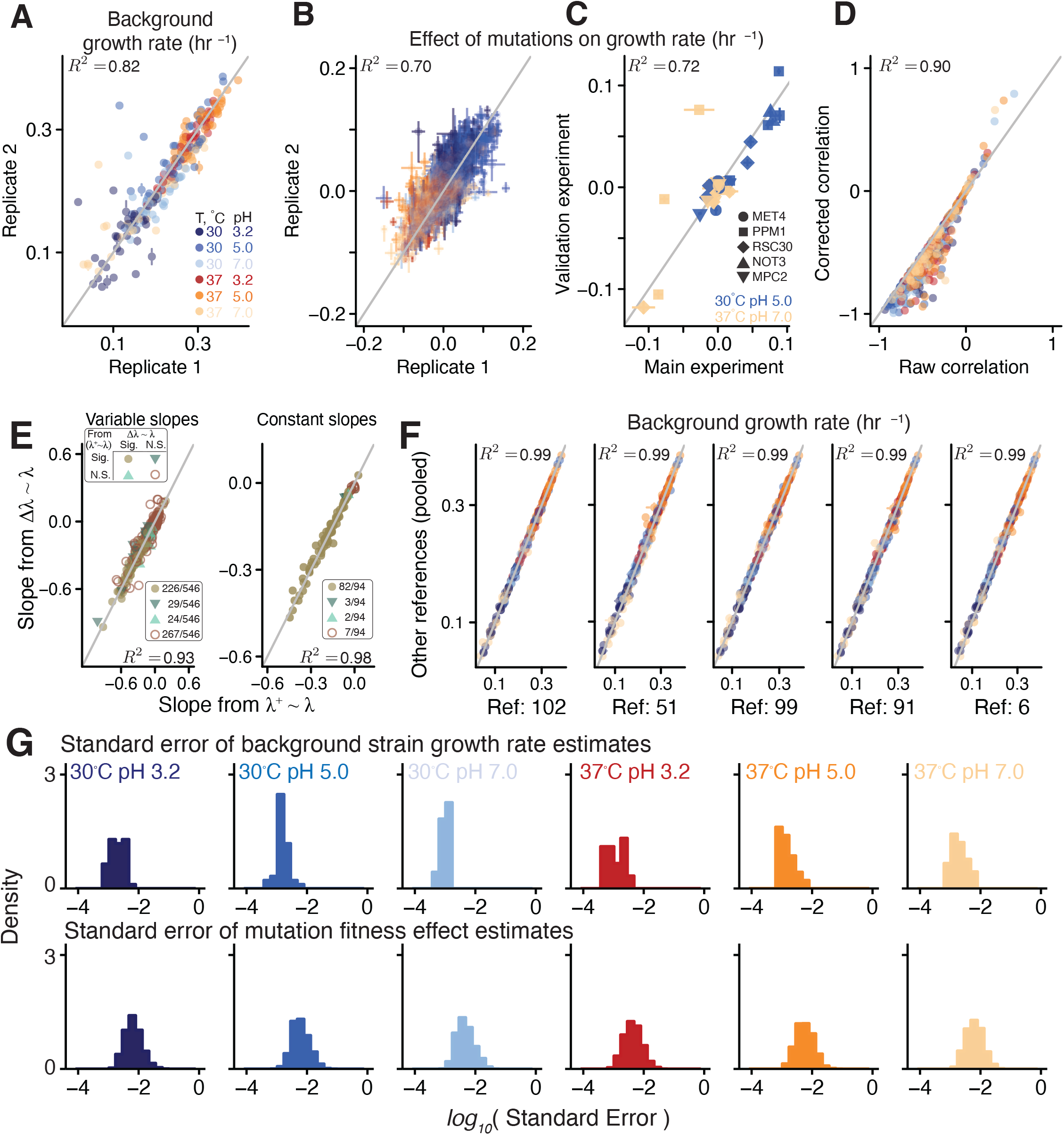
Data quality checks. **A**. Correlation between growth rates of background strains estimated in two biological replicates.**B**. As in panel A, but for the fitness effects of mutations. In both panels, error bars represent *±*1 standard error. **C**.Correlation between fitness effect estimates in the high-throughput RB-TnSeq experiment and the validation experiment (see Section 1.2.3). **D**. Raw and corrected estimates of the correlation coefficient between background growth rate and fitness effect for each mutation in each environment.**E**. Correlation between two approaches to estimate microscopic global epistasis slopes (see Section 1.3.7). Points are colored by the significance (*P <* 0.05) of the estimated slope. **F**. Correlation between background strain growth rates estimated from each neutral reference separately against the growth rate estimated from all other references pooled. Colors as in A. In all panels, grey line is the diagonal, *R*^2^ is reported for linear regression (*P <* 0.01 for all regressions). **G**. Distribution of the standard error of background strain growth rates (top) and mutation effects (bottom) in all environments.

**Figure S3.**
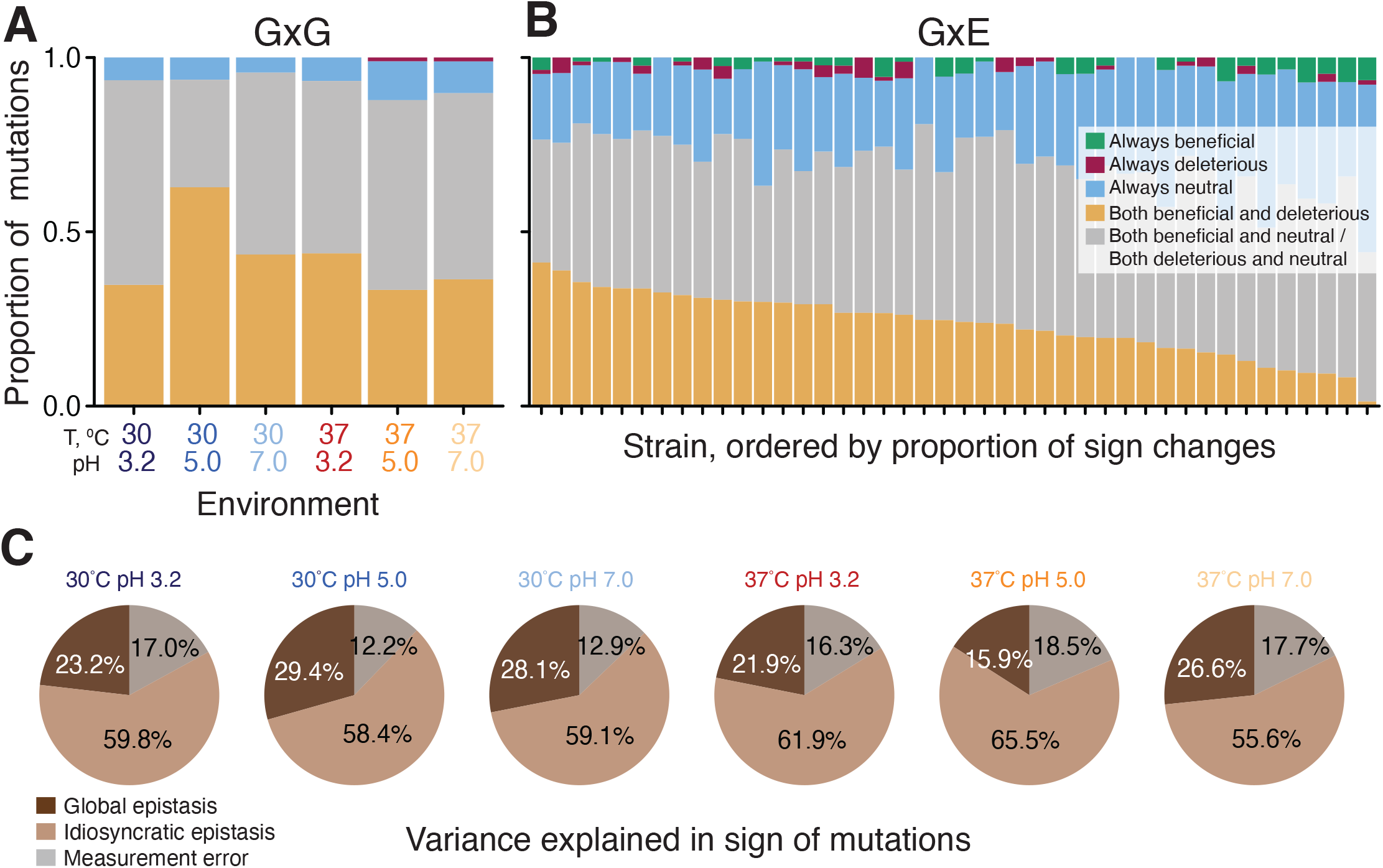
Variation in the sign of mutational effects across genetic backgrounds (G*×*G interactions) and across environments (G*×*E interactions). **A**. Proportions of mutations that do and do not change sign across background strains in each environment. **B**. Proportions of mutations that do and do not change sign across environments in each background strain. **C**. The proportion of variance in the observed sign of mutations in each environment explained by measurement noise, global and idiosyncratic epistasis (see Section 1.3.8 for details).

**Figure S4.**
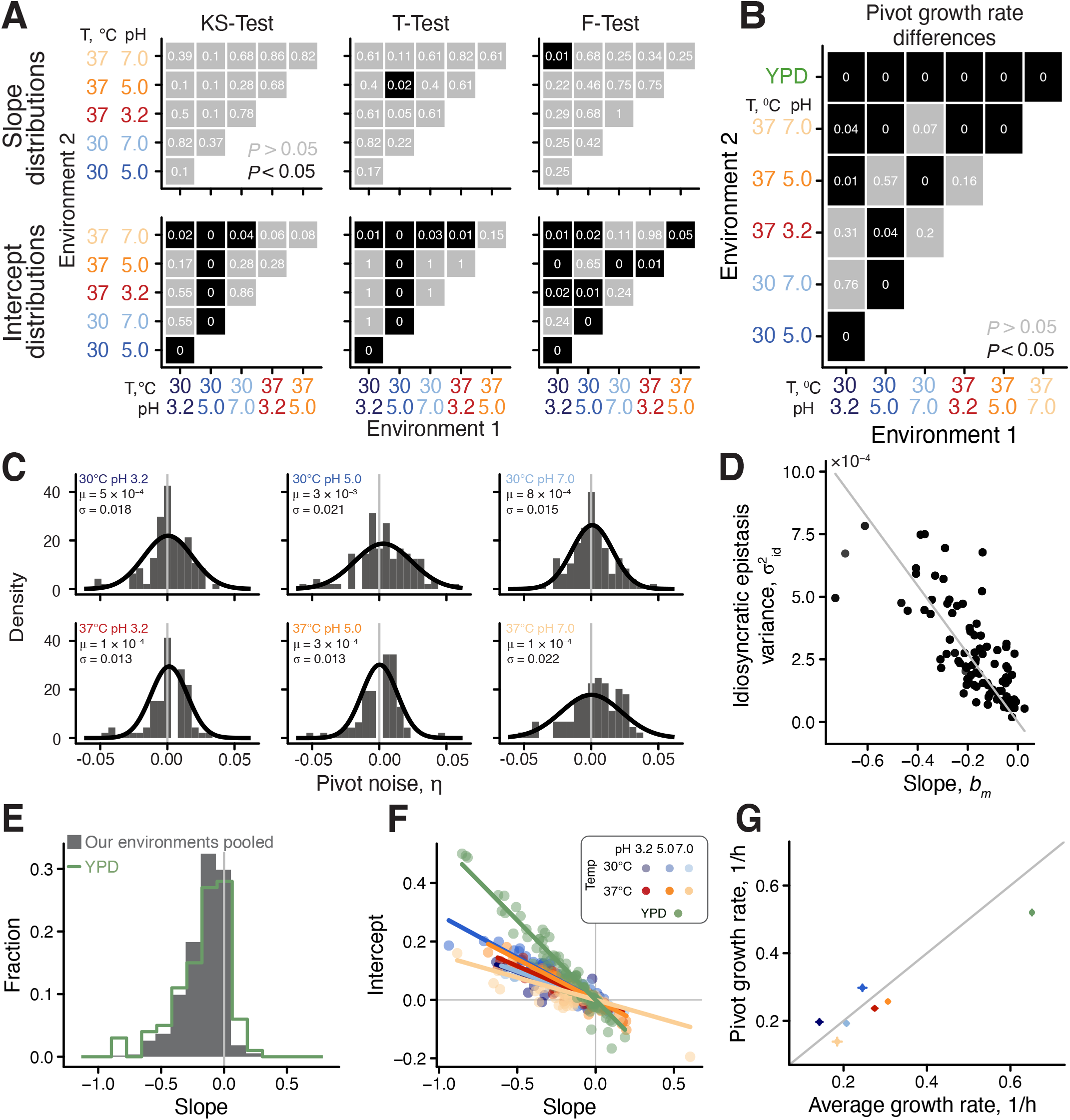
(Previous page) Properties of global epistasis models. **A**. Comparison of distributions of slopes (top) and intercepts (bottom) across environments using three metrics (see Section 1.3.7). **B**. Comparison of the pivot growth rates across environments. In both panels, the number in each tile is the *P* -value (after Benjamini-Hochberg correction), with values *P <* 0.01 indicated with ‘0’. Tiles with *P <* 0.05 are colored black. **C**. Distribution of the pivot noise term *η* in each environment. Mean and variance of the distribution are labelled in each panel, and the best fit normal distribution is overlayed. **D**. Relationship between global epistasis slope and the variance of idiosyncratic epistasis residuals from the fit of equation (2) in the main text. Grey line is best fit linear regression through the origin. **E**. Distribution of microscopic epistasis slopes in all our measured environments (pooled, grey) and in YPD (data from Ref. (*39*)). **F**. Slope-intercept relationship (as in Figure 2 in the main text), but including YPD data. **G**. The relationship between the average background strain growth rate and the pivot growth rate in each environment. Colors as in Panel F. Error bars are *±* 1 standard error.

**Figure S5.**
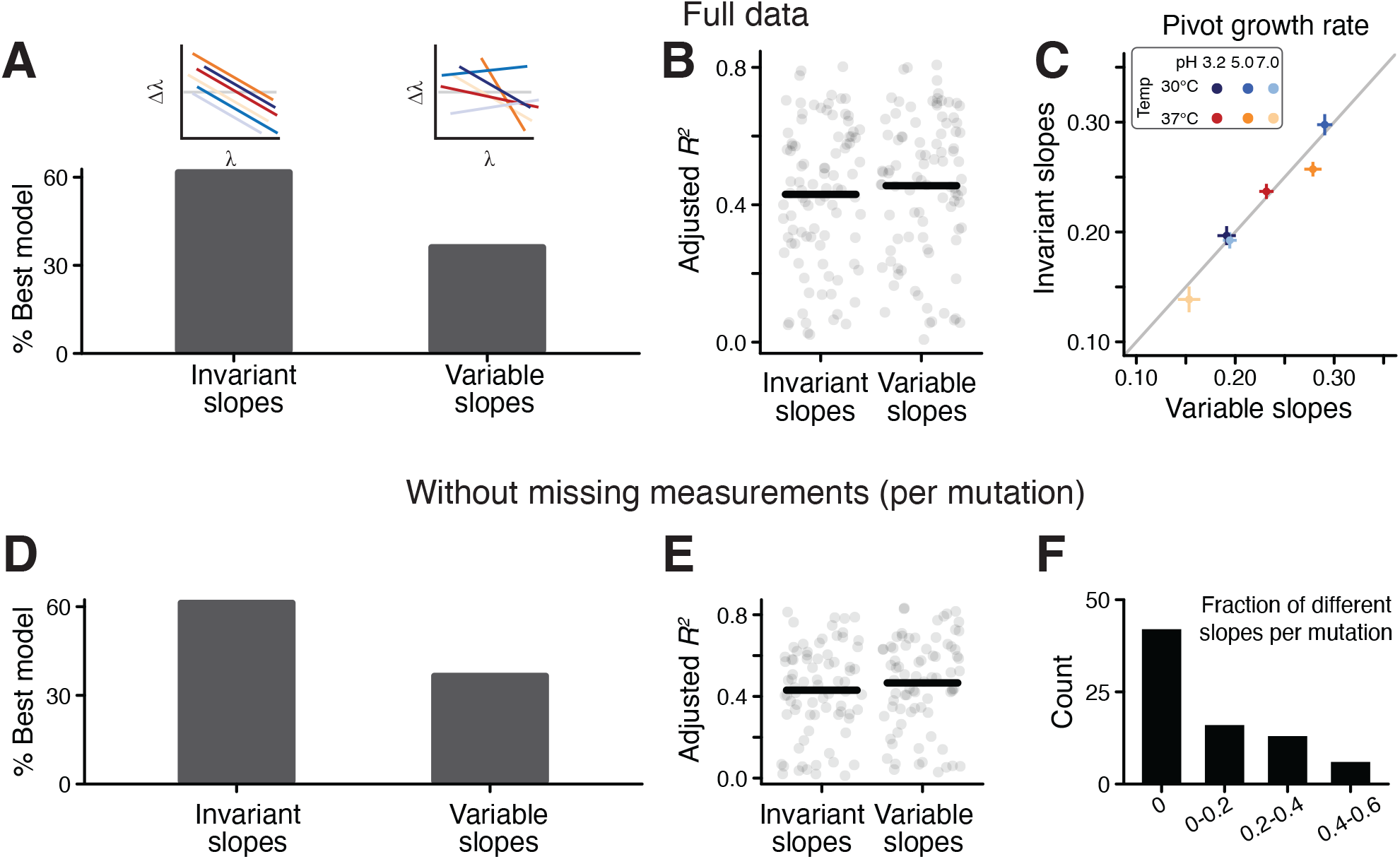
Variable and invariant slope models. **A**. Bar graph showing the percent of mutations best fit by the invariant slopes and variable slopes models (see Section 1.3.7) Schematic example of each model is shown on top. **B**. The adjusted *R*^2^ for the fits of both models for all mutations. **C**. Relationship between the pivot growth rates estimated from each model. Error bars are *±*1 standard error. **D, E**. As (A, B) but with a reduced data set with no missing measurements for each mutation (see Section 1.3.7). **F**. The fraction of significantly different slopes per mutation across environments in pairwise tests with the reduced data set (as Figure 3 in the main text).

**Figure S6.**
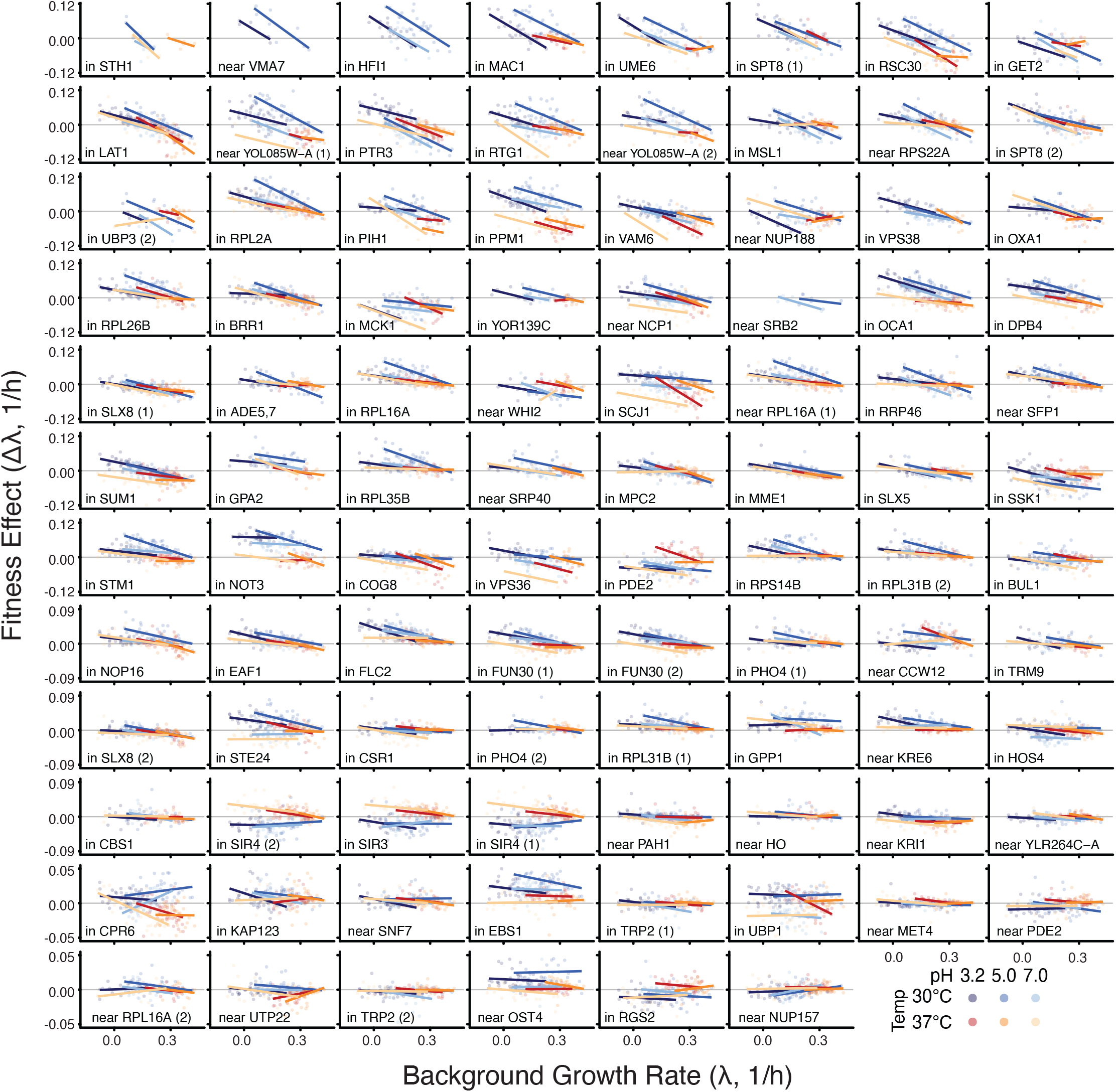
Global epistasis model for individual mutations. Same as Figure 3, but with data points. Mutations are ordered by slope, as in Figure 3. *y*-axis varies across rows.

**Figure S7.**
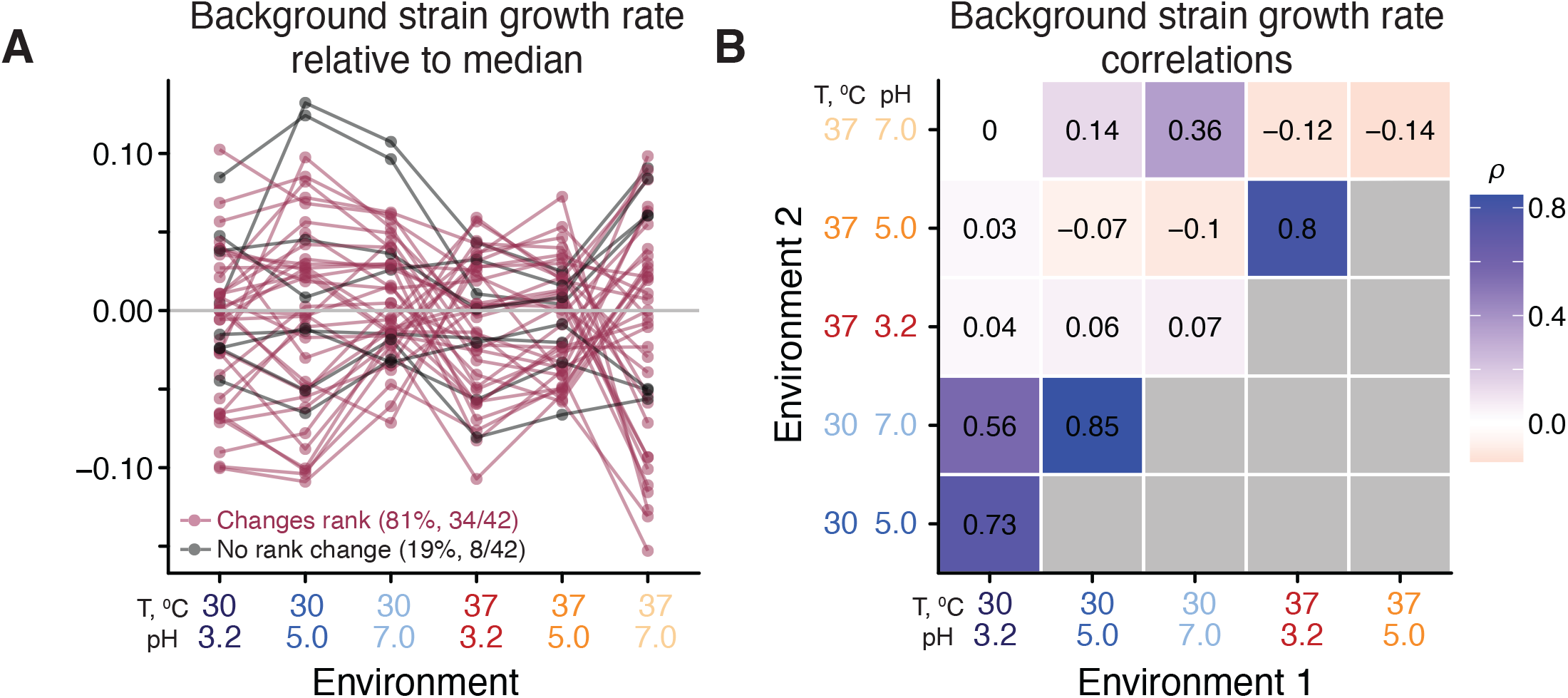
Reshuffling of background strain growth rates across environments. **A**. Strain growth rate relative to the median growth rate in each environment. Lines connect the same strain across environments and are colored maroon if the strain is on different sides of the median in different environments (see Section 1.3.6). **B**. The correlation of background strain growth rate across all pairs of environments.

**Figure S8.**
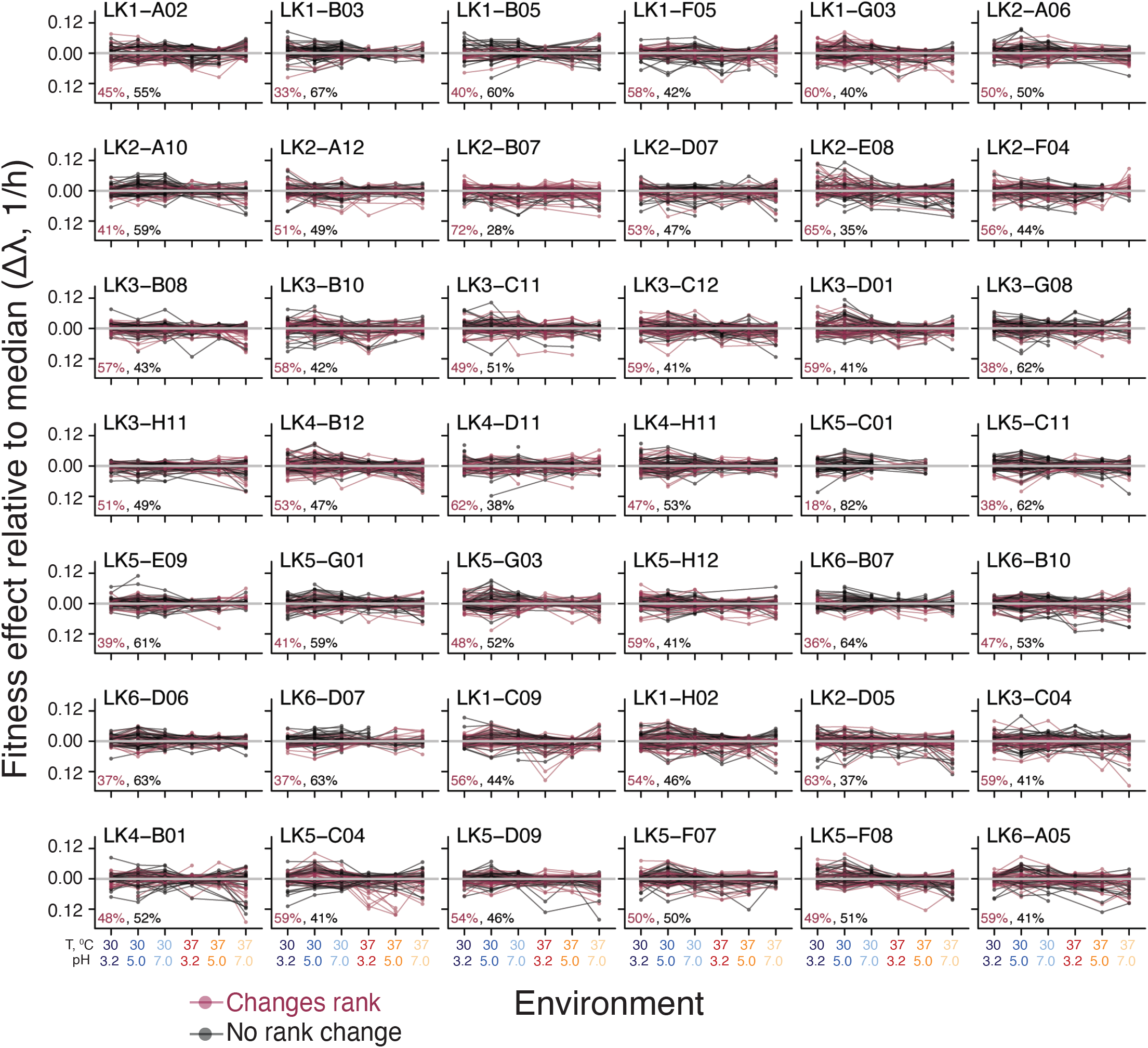
Reshuffling of the effects of mutations across environments. Each panel corresponds to a background strain and shows the effect of all mutations relative to the median in each environment. Lines connect the same mutation across environments and are colored maroon if the mutation is on different sides of the median in different environments (see Section 1.3.6).

**Figure S9.**
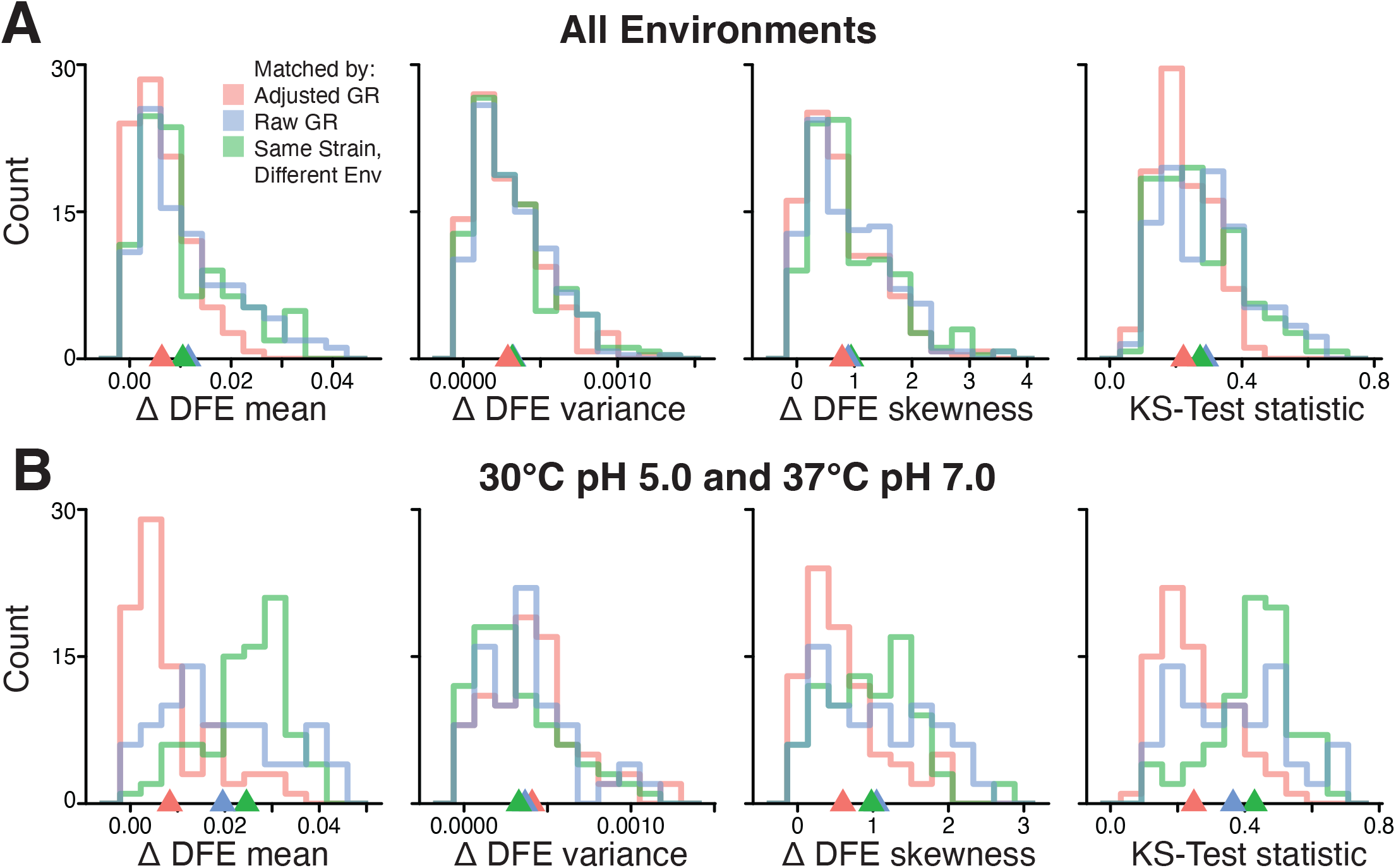
Distributions of DFEs similarity statistics. **A**. Distribution of four metrics of DFE similarity for all pairs of strains from any two different environments and matched either by their adjusted growth rate (red), raw growth rate (blue), or strain identity (green). See Section 1.3.9 for details. For all metrics, lower values mean more similar DFEs. Triangle shows the mean of the corresponding colored distribution. **B**. Same as A but with the strains sampled from the two most dissimilar environments, 30°C pH 5.0 and 37°C pH 7.0.

**Figure S10.**
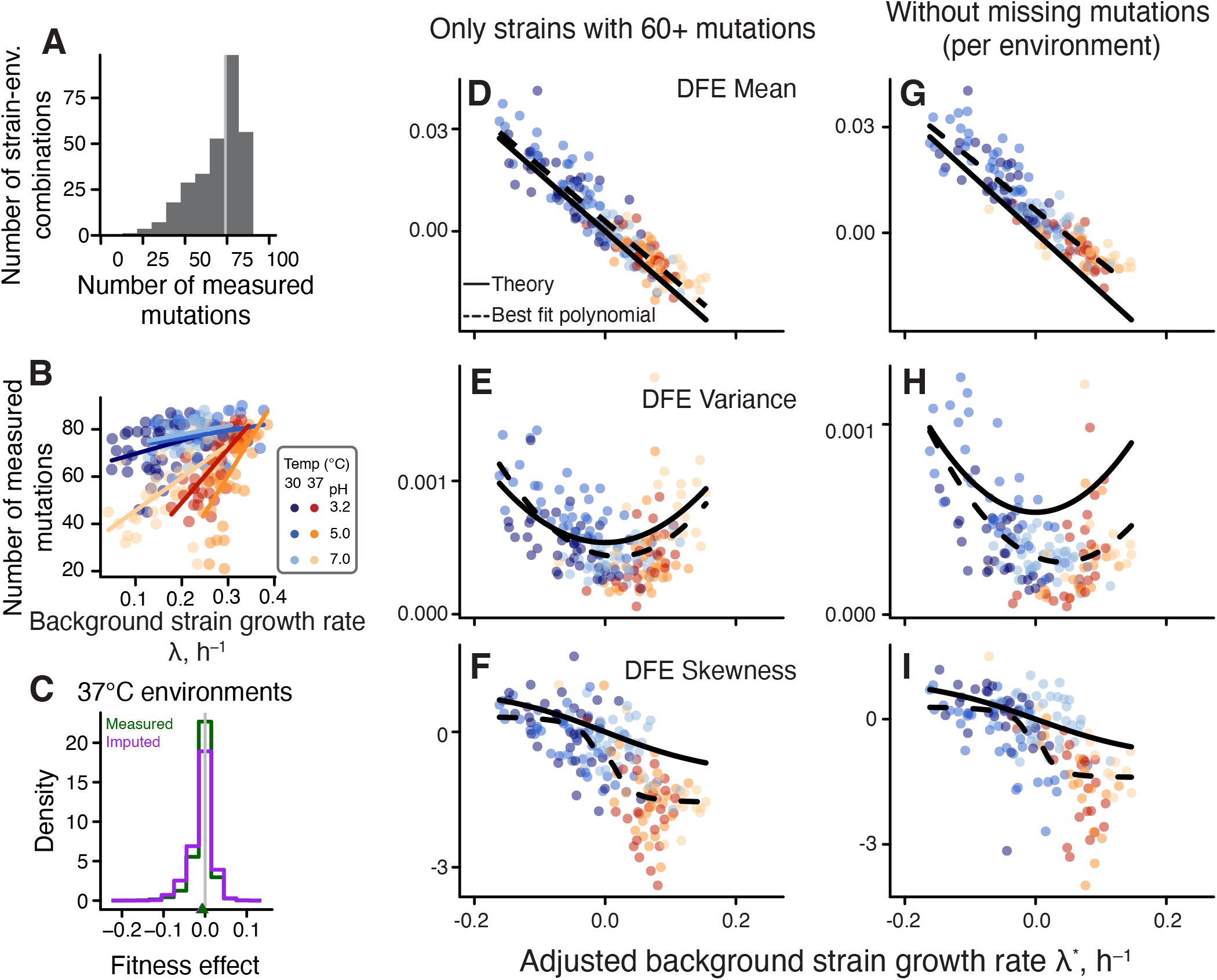
Robustness of the observed DFE variation with respect to missing measurements. **A**. Distribution of the number of mutations measured per DFE. **B**. Relationship between the number of mutations in each DFE and the background growth rate. Lines represent the best fit linear regression. **C**. Distribution of all measured mutation effects and the imputed missing mutation effects in 37°C environments (see Section 1.3.9). **D-F**. Same as Figure 4 but excluding all strains whose DFE contains less than 60 mutations. **G-I**. Same as Figure 4 but based on a reduced data set without missing measurements (see Section 1.3.9). Colors for D–I as in Panel B.

**Figure S11.**
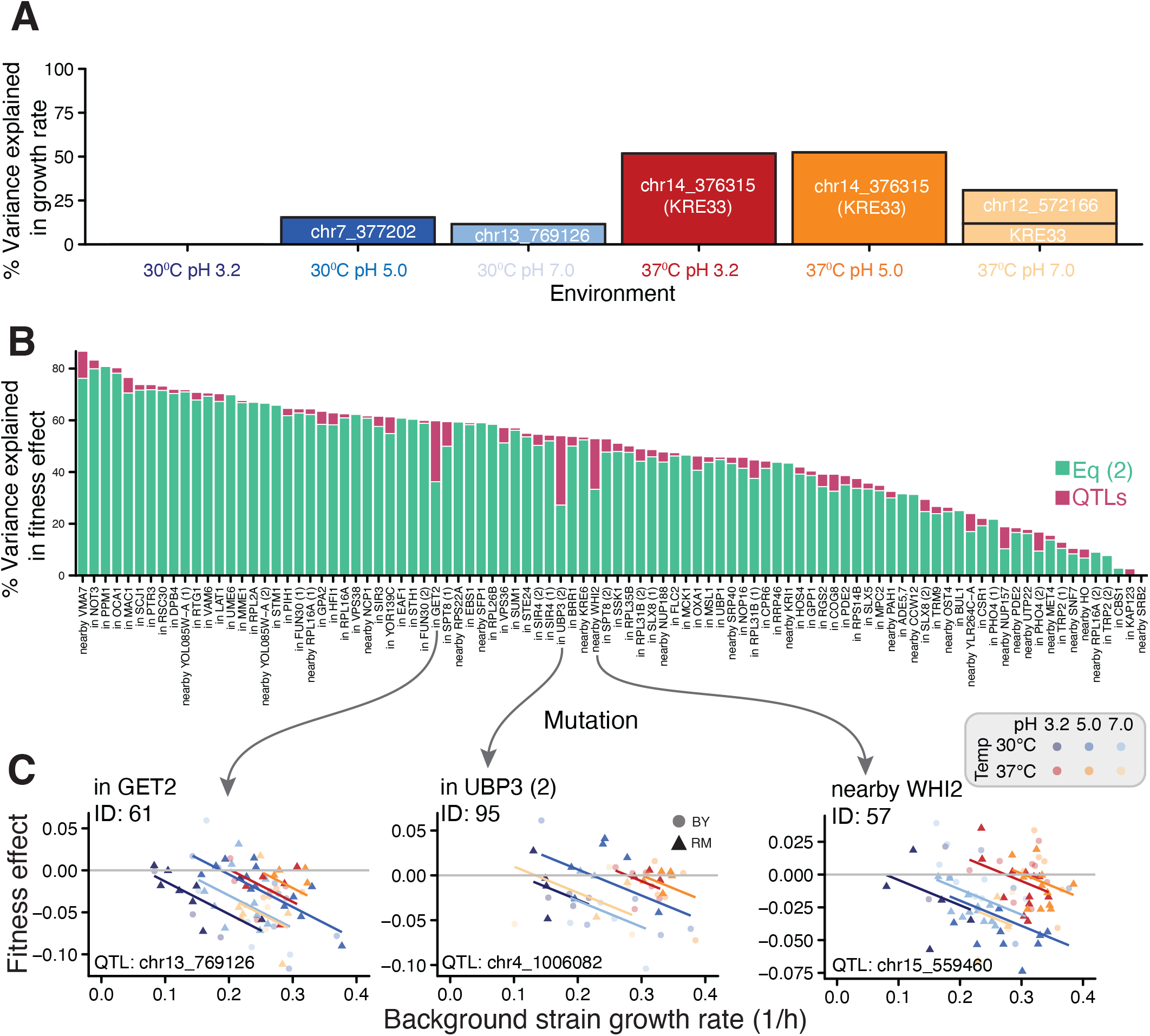
QTL analysis. **A**. Percent of variance in background strain growth rate explained by a model with 10 tested loci (see Section 1.3.10). Loci that explain a significant (*P <* 0.05) amount of variance are labelled, other loci are not shown. **B**. The percent variance in fitness effect of each mutation explained by 10 candidate loci combined (pink) above and beyond the variance explained by the generalized global epistasis equation (2) in the main text (teal). Mutations are ordered by the total explained variance. **C**. Three example mutations, with lines representing the best fit generalized global epistasis model, colored by environment. Point shape represents the allele, either BY (circles) or RM (triangles), at the locus explaining most variation for that mutation (locus indicated in the bottom left of each panel).

**Figure S12.**
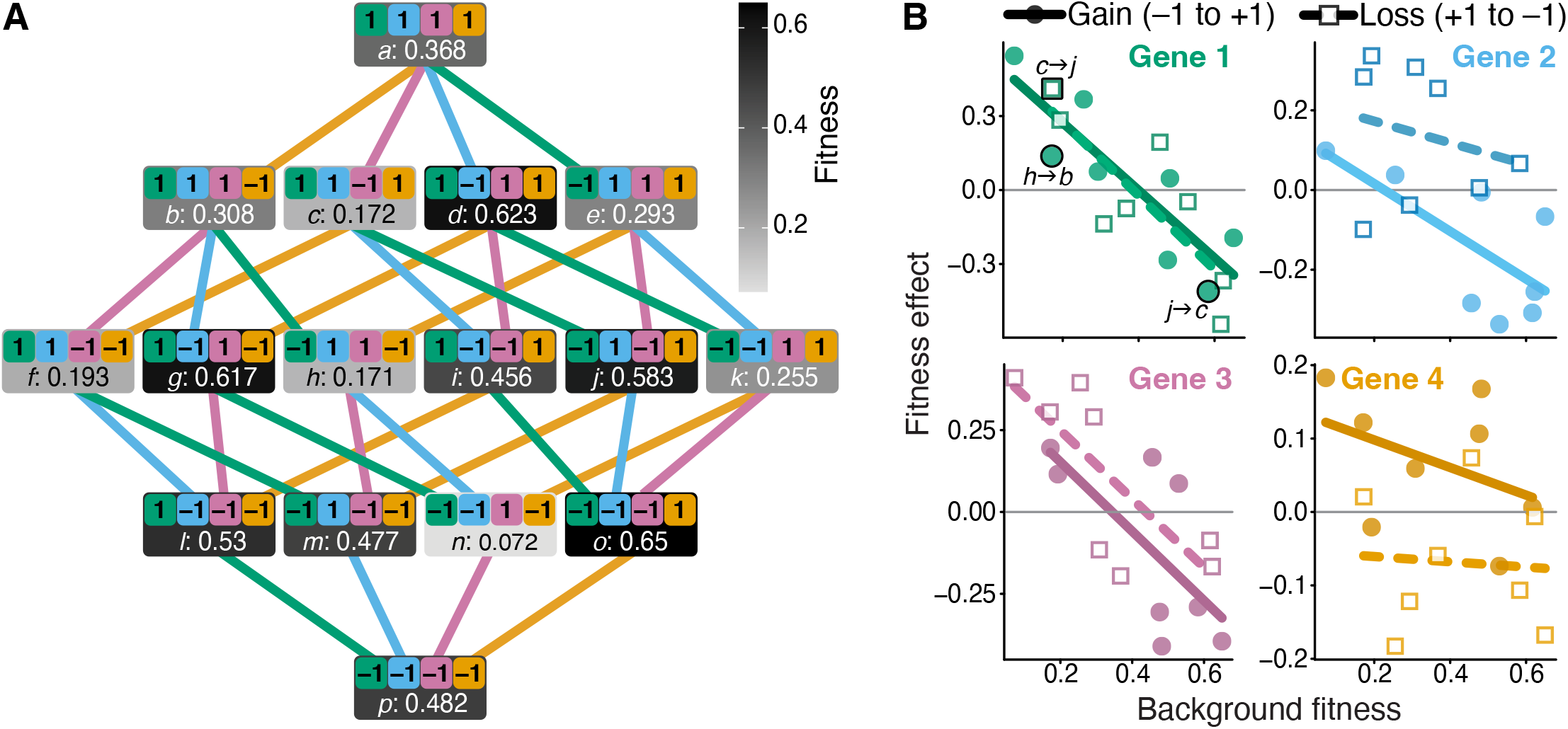
A binary 4-site genotype-to-fitness map that exhibits global epistasis. **A**. A graph representation of the genotype-to-fitness map. Each gray rectangle represents a genotype (sites are denoted by different colors, shade of gray represents fitness) and contains its shorthand letter notation and fitness value. Alleles +1 and *−*1 can be interpreted as the presence and absence of a functional gene. Lines connect genotypes separated by a single allele change (e.g., a single flip from +1 to *−*1 corresponding to a loss of one gene). Line color corresponds to the gene where the mutation occurs. **B**. Global epistasis plots for each of four sites (genes). Filled circles represent gene gains and empty squares represent gene losses. Solid and dashed lines show corresponding linear regressions. Some mutations mentioned in the text are outlined and labelled.

**Figure S13.**
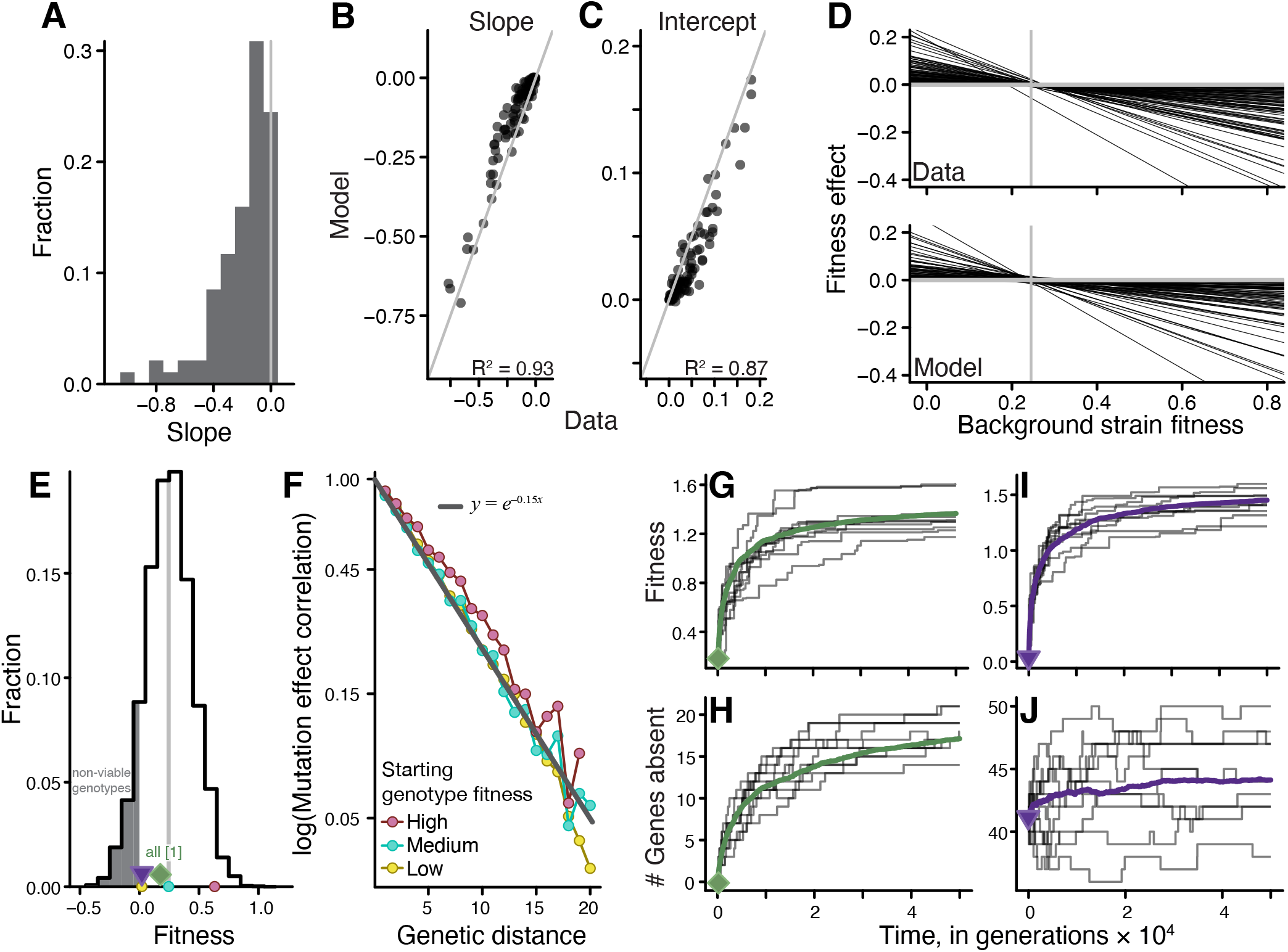
Connectedness (CN) model and evolutionary simulations. **A**. Distribution of global epistasis slopes in the concatenated data from our 30°C pH 5.0 environment (see Section 2.3.2). **B, C**. Correlation between the empirical global epistasis slopes and interceptes and those estimated from the CN model parameterized with the empirical slope distribution shown in panel A. Grey line is the identity line. **D**. “Bowtie” plot of all mutation regressions from the data (top) and the CN model (bottom). **E**. Fitness distribution of 5,000 random genotypes sampled from the CN model. Gray line indicates the pivot growth rate. Non-viable genotypes are shaded in gray. Colored circles indicate three genotypes in whose vicinity the mutation effect correlations are calculated (see panel F). Green rhombus (genotype with all intact genes) and purple triangle (low-fitness genotype) indicate the initial genotypes for which evolutionary simulations were carried out (see panels G–J). **F**. Mutation effect correlation *γ*_*d*_ plotted as a function of genetic distance *d* (see Section 2.3.2 for details) for three initial genotypes (see panel E). Gray line is the best-fit exponential. **G**. Fitness in Wright-Fisher simulations with *NU* = 0.1 (Section 2.3.2), initiated with a genotype with all intact genes (see panel E). Thick green line is the mean over 50 replicates, with 10 random individual trajectories shown in gray. **H**. Count of intact genes in these simulations. **I, J**. Same as G, H, but with the initial low-fitness genotype (panel E).

## 4 Supplementary tables

**Table S1.** Statistics of slope and intercept distributions.

| Environment |  | Slopes |  |  | Intercepts |  |  |
| --- | --- | --- | --- | --- | --- | --- | --- |
| Temp, °C | pH | Mean | Stdev | Skew | Mean | Stdev | Skew |
| 30 | 3.2 | −0.160 | 0.143 | −1.40 | 0.0326 | 0.0313 | 0.824 |
| 30 | 5.0 | −0.214 | 0.186 | −0.922 | 0.0640 | 0.0556 | 0.404 |
| 30 | 7.0 | −0.161 | 0.157 | −0.824 | 0.0316 | 0.0337 | 0.230 |
| 37 | 3.2 | −0.136 | 0.151 | −1.09 | 0.0311 | 0.0378 | 0.581 |
| 37 | 5.0 | −0.121 | 0.157 | −0.839 | 0.0292 | 0.0498 | 0.324 |
| 37 | 7.0 | −0.137 | 0.189 | −0.731 | 0.0169 | 0.0401 | −0.790 |

**Table S2.** Antibiotic concentrations used in this study, in µg/ml.

| **Antibiotic** | ***E. coli*** | ***S. cerevisiae*** |
| --- | --- | --- |
| Kanamycin (Kan) | 40 | N/A |
| Ampicilin (Amp) | 100 | 100 |
| Nourseothricin (Nat) | 20 | 20 |
| Hygromycin (Hyg) | 200 | 300 |

**Table S3.** Best fit trends in DFE moments. “Pwr” refers to the power of to each the predictor (background growth rate) is raised in each term of the polynomial (see 1.4). “Factor” is the scaling factor by which all entries in the row are multiplied. “Reduced data 1” is the data set with only DFEs made of *>* 60 mutations. “Reduced data 2” is the data set with no missing mutations per environment. “Est” refers to the best-fit parameter value.

| Moment | Pwr | Factor | Theory | Full data |  | Reduced data 1 |  | Reduced data 2 |  |
| --- | --- | --- | --- | --- | --- | --- | --- | --- | --- |
|  |  |  |  | Est. | 95% CI | Est. | 95% CI | Est. | 95% CI |
| Mean | 0 | $10^{-3}$ | 0 | 2.2 | [1.5, 2.9] | 2.9 | [2.2, 3.6] | 6.4 | [5.7, 7.2] |
| | 1 | $(-10^{-1})$ | 1.7 | 1.7 | [1.8, 1.7] | 1.6 | [1.7, 1.5] | 1.5 | [1.6, 1.4] |
| Var | 0 | $10^{-4}$ | 5.4 | 4.6 | [4.3, 5.0] | 4.4 | [4.0, 4.9] | 3.0 | [2.6, 3.4] |
| | 1 | $10^{-4}$ | 0 | 2.0 | [-2.5, 6.5] | -7.8 | [-12, -3.2] | -13 | [-18, -9] |
| | 2 | $10^{-2}$ | 1.7 | 2.4 | [2.1, 2.8] | 2.1 | [1.6, 2.7] | 1.7 | [1.2, 2.2] |
| Skew | 0 | $(-1)$ | 0 | 0.39 | [0.52, 0.27] | 0.41 | [0.56, 0.26] | 0.27 | [0.47, 0.07] |
| | 1 | $10^{-2}$ | -3.1 | -4.6 | [-5.7, -3.4] | -5.4 | [-6.9, 3.9] | -2.5 | [-3.7, -1.5] |
| | 2 | $10^{-1}$ | 0 | 0.02 | [-1.5, 1.5] | -0.7 | [-2.4, 1.0] | -0.13 | [-1.6, 1.3] |
|  | 3 | 1 | -1.12 | -1.21 | [-2.10, -0.33] | -1.52 | [-3.07, 0.03] | -1.03 | [-2.31, 0.25] |

**Table S4.**
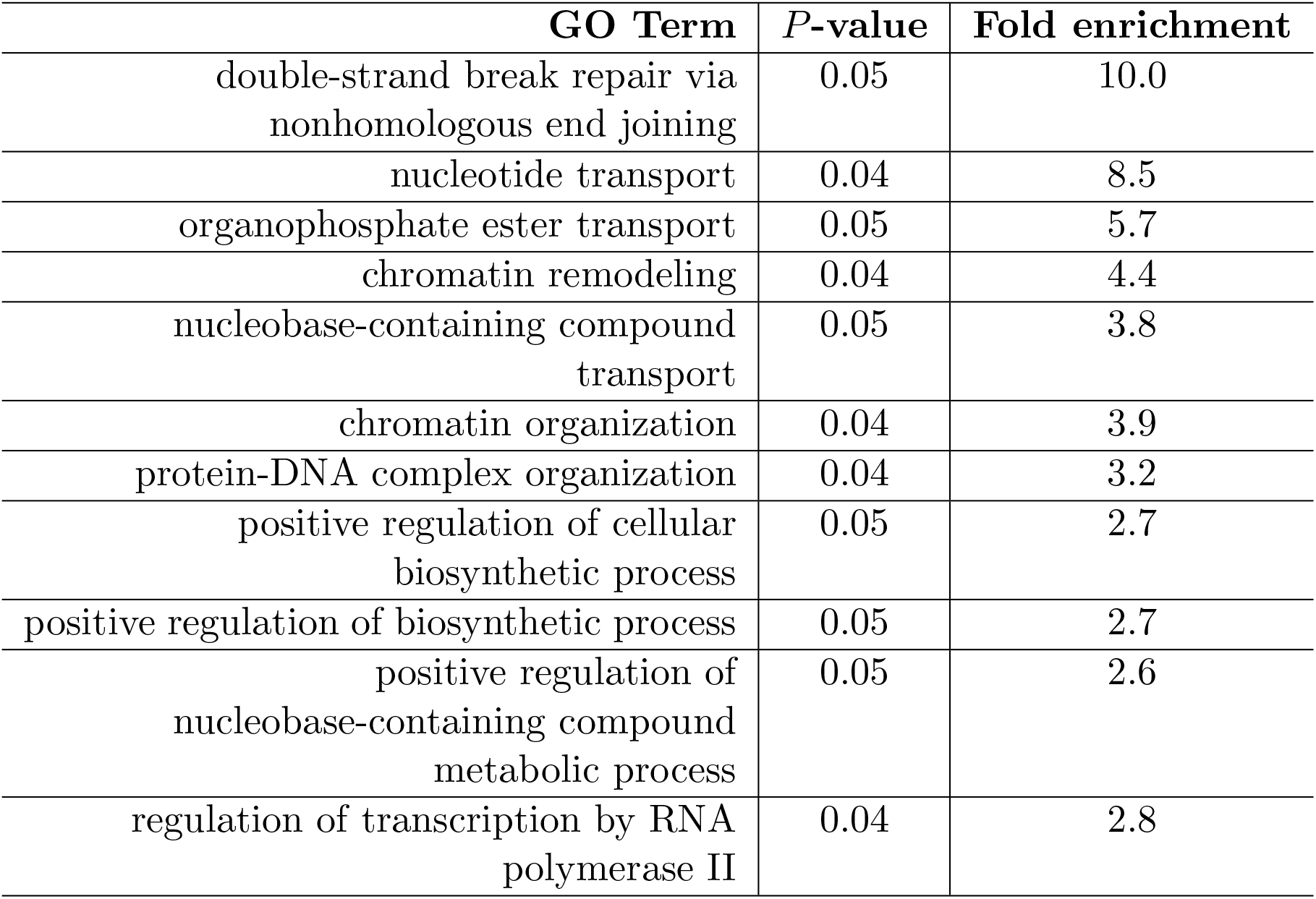
GO term enrichment among all mutations in this study. *P* -value is reported after the Benjamini-Hochberg correction.

## Footnotes

1 This was done to avoid testing correlated effects of linked QTLs.

2 Unlike the environments in the present work, the SC 37°C medium in Ref. (*34*) was not buffered.

3 According to the Reddy-Desai model, the pivot growth rate equals the average fitness of all genotypes (*44*). The fact that the average growth rate of our background strains correlates with the pivot growth rate (Figure S4G) suggests that our background genotypes are not particularly exceptional.

4 Our focus in this paper has been on the deterministic fitness-correlated trends in the shape of the DFE, but Figure 4D shows that there is also a considerable amount of variation around these trends on which second-order selection could potentially act. Furthermore, Johnson and Desai find that clones sampled from the same population at the same time point sometimes have significantly different DFEs, which further supports the existence of meaningful DFE variation on which second-order selection could act.

5 In this model, the two alleles +1 and *−*1 are arbitrary labels and therefore completely equivalent, which of course may not be the case in real biological systems. Moreover, this model assumes that none of the genes are essential, so that, for example, the genotype with all *−*1 alleles is viable. Note that in our experiment we also consider only non-lethal tn-insertion mutations (see Section 1.1.2).

6 In principle, the CN model can generate additive landscapes too, but these should be exceedingly rare.

